# Selective G protein signaling driven by Substance P-Neurokinin Receptor structural dynamics

**DOI:** 10.1101/2021.05.16.444192

**Authors:** Julian A. Harris, Bryan Faust, Arisbel B. Gondin, Marc André Dämgen, Carl-Mikael Suomivuori, Nicholas A. Veldhuis, Yifan Cheng, Ron O. Dror, David M. Thal, Aashish Manglik

## Abstract

The neuropeptide Substance P (SP) is important in pain and inflammation. SP activates the neurokinin-1 receptor (NK1R) to signal via G_q_and G_s_ proteins. Neurokinin A also activates NK1R, but leads to selective G_q_ signaling. How two stimuli yield distinct G-protein signaling at the same G-protein-coupled-receptor remains unclear. We determined cryo-EM structures of active NK1R bound to SP or the G_q_-biased peptide SP6-11. Peptide interactions deep within NK1R are critical for receptor activation. Conversely, interactions between SP and NK1R extracellular loops are required for potent G_s_ signaling but not G_q_ signaling. Molecular dynamics simulations showed that these superficial contacts restrict SP flexibility deep in the NK1R pocket. SP6-11, which lacks these interactions, is dynamic while bound to NK1R. Structural dynamics of NK1R agonists therefore depend on interactions with the receptor extracellular loops and regulate G-protein signaling selectivity. Similar interactions between other neuropeptides and their cognate receptors may tune intracellular signaling.

## Introduction

Substance P (SP) is a peptide with incredibly diverse roles in animal physiology. Like other neuropeptides, SP exerts long-lasting regulation of synaptic neurotransmission by activating its cognate G protein-coupled receptor (GPCR), the neurokinin 1 receptor (NK1R). SP action in the nervous system is important in pain, mood, respiration, and nausea^1–3^. Action of SP in other tissues is associated with inflammation or smooth muscle contraction^1,2^. Extensive studies suggest that inhibition of SP activity by NK1R antagonists might lead to effective treatments for pain, inflammation, and mood disorders^4,5^, although the only clinical success to date has been for treatment of chemotherapy-induced nausea and vomiting^6,7^

The NK1R is endogenously activated by SP and another neuropeptide, neurokinin A (NKA). Both SP and NKA belong to the larger family of tachykinin neuropeptides that share a common C-terminal ‘F(V/F)GLM-NH_2_’ consensus sequence, which is required for their activity at any of the three neurokinin receptors. The more divergent N-terminal region of tachykinin peptides has previously been implicated in dictating which neurokinin receptor a tachykinin prefers^8,9^. Like other neuropeptides, tachykinin function follows the “message-address” framework, in which two distinct portions of a peptide encode either the efficacy (message) or receptor selectivity (address)^10–12^. Following this framework, NKA was initially described as specific for the neurokinin 2 receptor (NK2R)^8,13^. However, both SP and NKA activate NK1R in cell lines and in various physiological settings^14–18^.

Intriguingly, activation of NK1R by SP or NKA induces distinct cellular responses and, in certain tissues, distinct physiological outcomes^16,19–21^. SP increases both inositol phosphate (IP) and cAMP second messengers downstream of G_q_ and G_s_ signaling pathways, respectively^17,18,22^. By contrast, NKA signals potently via G_q_ but has decreased G_s_ stimulatory activity^17,18^. Molecular pharmacology studies revealed that SP binding to NK1R is distinct from NKA binding^18,23,24^. A common model proposed by these studies is that NK1R exists in two distinct active conformations: an SP-selective state and a general-tachykinin state that binds both SP and NKA^18,24^. Mutations can alter the relative proportion of these two states, yielding changes in the measured affinities for SP, NKA and related tachykinins^18,24^. Intriguingly, these mutations also dramatically affect the ability of NK1R to signal via G_q_ or G_s_, suggesting that these distinct active conformations are coupled to distinct signaling outcomes^18^.

The ability of two agonists to induce distinct intracellular signaling cascades downstream of a single GPCR is well established. However, how two endogenous stimuli yield distinct G protein coupling preference at the same receptor remains unclear at the biochemical and structural level. Here, we combine structural biology with molecular dynamics simulations and cellular signaling studies to decipher the molecular basis of agonist-dependent G protein-selective signaling at the NK1R.

### Structure of the Substance P-NK1R-miniG_s/q70_ complex

To enable structure determination of active human NK1R without thermostabilizing mutations or truncations, we generated a construct with the engineered Gα subunit miniG_s/q70_ fused to the C-terminus of the receptor^25^. The miniG_s/q70_ protein presents the GPCR-interacting α5 helix of Gα_q_ on an engineered Gα_s_ protein stabilized in the active conformation and with complete truncation of the Gα alpha-helical domain. This strategy improved the biochemical stability of Substance P (SP)-bound receptor compared to full-length NK1R alone (Supplementary Fig. 1). Purified SP-bound NK1R-miniG_s/q70_ fusion protein was mixed with excess Gβγ and nanobody 35 (Nb35)^26^ for structure determination by cryogenic-electron microscopy (cryo-EM) (Supplementary Fig. 1).

We determined a cryo-EM structure of the SP-NK1 R-G_s/q70_ complex at a global resolution of 3.0 Å (Fig. 1A and Supplementary Fig. 2). As is common for many GPCR-G protein complex structures, our initial maps yielded poor resolution for SP, the orthosteric binding pocket, and the extracellular loops. To improve reconstruction in these regions, we performed iterative focused refinements using a mask encompassing only the upper transmembrane region of the 7TM bundle. The resulting improved maps enabled an atomic model for all subunits of the complex and the SP peptide (Fig. 1B, Supplementary Fig. 2, Supplementary Fig. 3).

**Figure 1.**
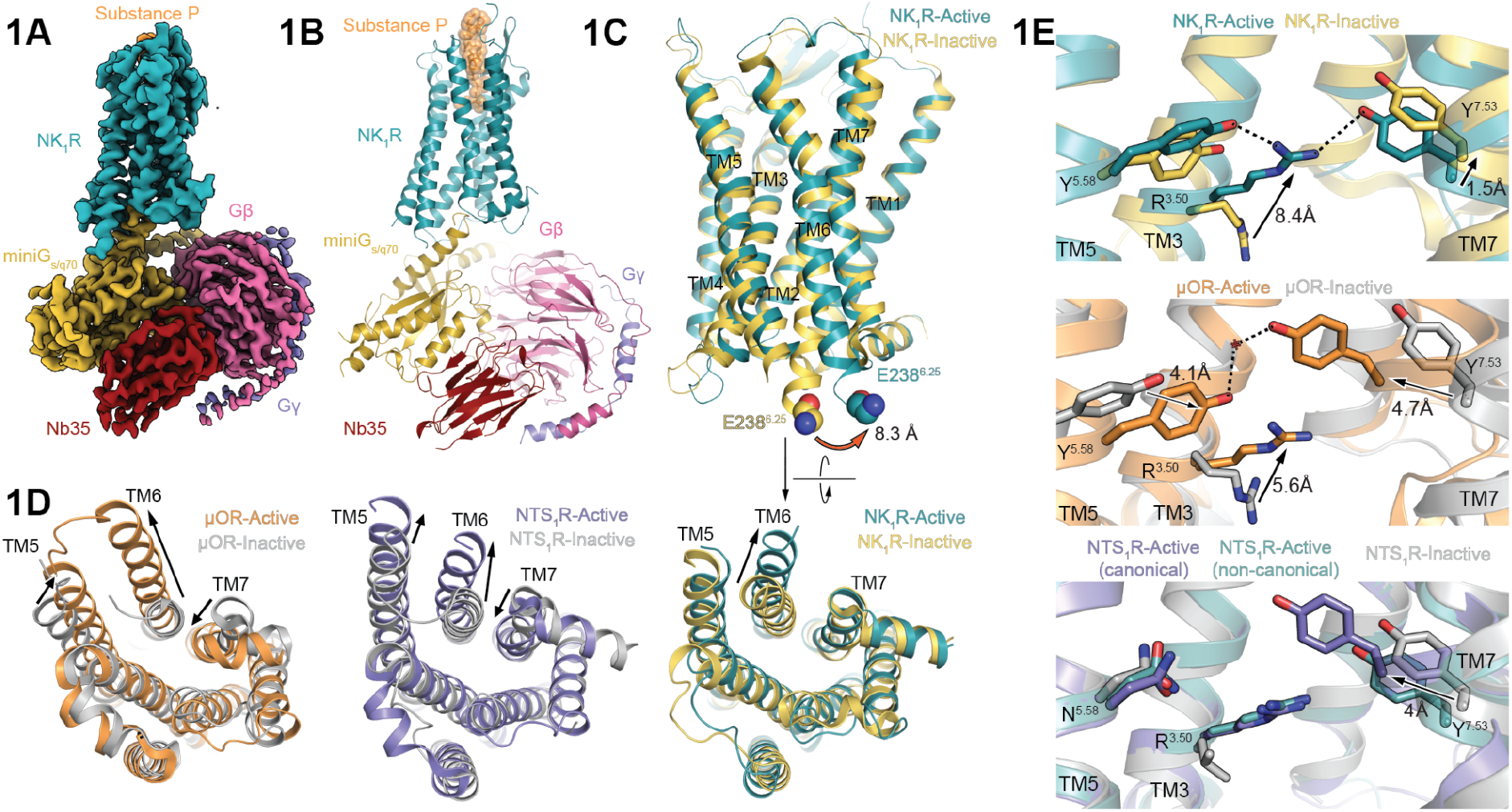
Cryo-EM structure of active NK1R bound to Substance P. **(A)** Unsharpened cryo-EM map of Substance P-bound NK1R-miniG_s/q70_-Nb35 complex. **(B)** Ribbon diagram of NK_1_R-miniG_s/q70_-Nb35 complex. Substance P is shown as orange spheres. **(C)** Alignment of active NK_1_R with inactive, antagonist-bound NK1 R (PDB: 6HLP^38^) shows 8.3 Å outward displacement of TM6. **(D)** Comparison of active NK_1_R to other active-state GPCRs shows minimal inward movement of TM7 upon activation. Activation-dependent inward movement of TM7 for two class A neuropeptide GPCRs is shown for comparison: μ-opioid (μOR, active PDB: 5C1M^29^, inactive PDB: 4DKL^58^) and neurotensin 1 (NTS_1_R, active PDB: 6OS9^35^, inactive PDB: 4BUO^59^). **(E)** The active-state NK1 R NPxxY motif shows a similar conformation to the ‘non-canonical’ active-state NTS_1_R conformation (PDB: 6OSA^35^). Inward movement of Y^7.53^ and TM7 for canonically active μOR and NTS_1_R is shown for comparison.

SP-activated NK1R is in a distinct active conformation when compared to other class A GPCRs. Like other GPCRs, active NK1R displays an 8.3 Å movement of transmembrane helix 6 (TM6) away from the 7TM helical bundle, enabling insertion of the C-terminal α5-helix of the miniG_s/q70_ protein (Fig. 1C). This movement is associated with other conserved changes in class A GPCR activation, including displacement of the W^6.48^ ‘toggle-switch’ (superscripts denote Ballesteros-Weinstein numbering^27^), rearrangement of the ‘P^5.50^I^3.40^F^6.44^’ connector motif, and movement of the ‘D^3.49^R^3.50^Y^3.51^’ motif (Supplementary Fig. 4)^28^. These conformational changes link ligand binding in the orthosteric site to the intracellular G protein coupling site and facilitate G protein binding.

By contrast, the conserved ‘N^7.49^P^7.50^xxY^7.53^’ motif in the SP-NK1R-miniG_s/q70_ structure remains in an inactive conformation. A hallmark of class A GPCR activation is inward movement of TM7 into the helical core^28^ (Fig. 1D). This allows Y^7.53^of the NPxxY motif to engage in an extended water-mediated hydrogen bonding-network with other residues on the cytoplasmic face of TM3 and TM5, as observed for the active μ opioid receptor^29^ (Fig. 1E). This inward movement of Y^7.53^ is not observed in the SP-NK1R-miniG_s/q70_ structure and TM7 remains in a conformation that closely resembles the inactive-state (Fig. 1E). Importantly, other structures of GPCRs solved in complex with G_q/11_ family G proteins, miniG_s_, and miniG_s/q70_ proteins show canonical inward movement of TM7 upon receptor activation^30–34^.

The unique active-state of SP-bound NK1R resembles a previously determined structure of the neurotensin 1 receptor (NTS_1_R) bound to the cAMP inhibitory G protein G_i_^35,36^. Two active-state conformations of NTS_1_R bound to G_i_ have previously been observed: a canonical state with inward movement of TM7 and a ‘non-canonical’ state without TM7 rearrangement^35^ (Fig. 1E). Although the seven transmembrane domain of NK1R bound to miniG_s/q70_ is in a similar conformation to the ‘non-canonical’ NTS_1_R conformation, we do not observe the 45° rotation of the G protein observed for ‘non-canonical, NTS_1_R (Supplementary Fig. 4). While there are important caveats to our interpretation of the interactions between NK1R and the engineered miniG_s/q70_ protein, we surmise that fully active NK1R bound to the C-terminus of G_q_ exists in a unique conformation compared to most class A GPCRs.

Unlike the majority (98%) of class A GPCRs, NK1R possesses a glutamate residue at the highly conserved D^2.50^ position^37,38^. In inactive-state class A GPCRs, this canonical D^2.50^ residue participates in an extended, water-mediated hydrogen-bonding network between TM helices 2,3,6 and 7. For most GPCRs, activation is coupled with an inward movement of TM7 driven by a direct interaction between D^2.50^ and N^7.49^ (Supplementary Fig. 4) of the NPxxY motif. By contrast, E78^2.50^ forms a direct interaction with N301^7.49^ in the NK1R inactive-state. We speculate that the stable and direct E^2.50^-N^7.49^ interaction in the NK1R inactive-state disfavors inward TM7 motion during activation and contributes to the ‘non-canonical’ active-state. Indeed, prior work has shown that disrupting the E^2.50^-N^7.49^ interaction with mutagenesis selectively diminishes G_s_ signaling but does not affect G_q_ signaling^39^, suggesting that the ‘non-canonical’ NK1R active conformation is important for robust G_s_ and G_q_ signaling downstream of NK1R activation.

### Molecular Recognition of Substance P by NK1R

SP binds with an expansive interface stretching from a deeply-buried 7TM pocket to the distal portions of the NK1R ECL2 and N-terminus (Fig. 2A). We observed clearly resolved cryo-EM density for SP C-terminal residues 6-11, enabling us to unambiguously model this portion of the peptide (Fig. 2A). The N-terminal portion of SP, including residues 1-5, interact primarily with the extracellular loop 2 (ECL2) and the N-terminus of NK1R. The density for these residues is less well resolved, but we were able to confidently place all mainchain atoms and all side chains with the exception of R1 and K3.

**Figure 2.**
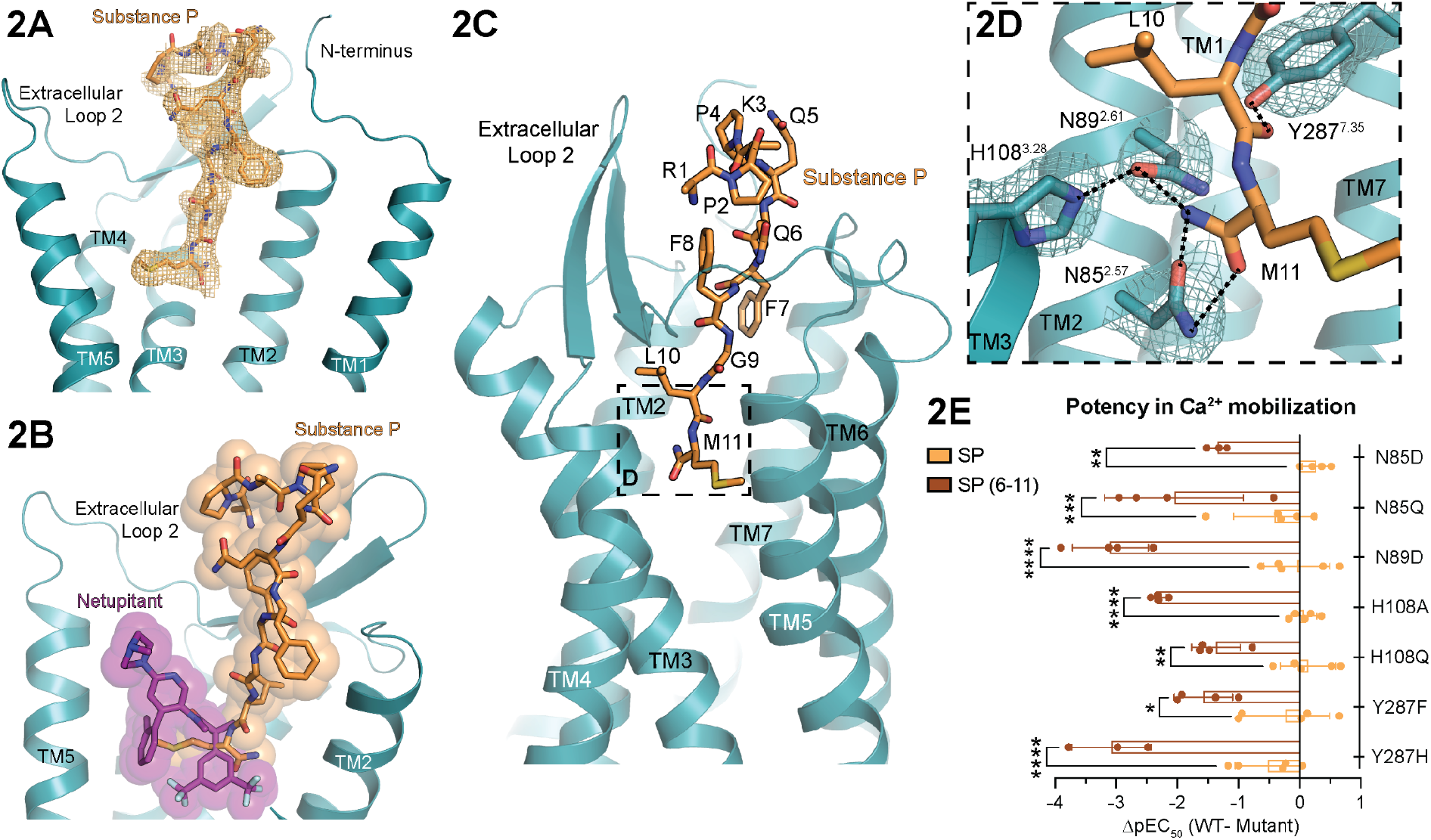
Molecular recognition of Substance P by NK1R. **(A)** Sharpened cryo-EM density map for Substance P (SP) in the NK1R binding pocket shown as orange mesh and contoured at a distance of 1.85 Å from the placement of Substance P atoms. **(B)** Overlay of Substance P and netupitant binding sites in NK1 R orthosteric site. **(C)** Substance P forms an extensive interaction interface with NK1 R, reaching from the deep orthosteric pocket to the distal extracellular regions. **(D)** The C-terminally amidated methionine of Substance P (M11) forms an extended hydrogen-bonding network with NK1R. Sharpened cryo-EM density map for sidechains is contoured 1.85 Å away from modeled atoms. **(E)** Truncated SP6-11 is sensitive to mutations in the deep orthosteric pocket, highlighting the importance of the extended hydrogen-bond network for Substance P recognition. Bar graphs represent mean ΔpEC_50_ (WT - Mutant) ± s.e.m. from n ≥ 3 independently fit biological replicates. Statistical significance between SP and SP6-11 ΔpEC_50_ for each mutant is compared in a two-way analysis of variance (ANOVA) with Šídák’s multiple comparison-corrected post hoc test, (* = p ≤ 0.033, ** = p ≤ 0.002, *** = p ≤ 0.0002, **** = p ≤ 0.0001). Full quantitative parameters from this experiment are listed in Supplementary Table 3.

SP binds to NK1R in a distinct manner compared to other neuropeptides at their cognate receptors. We compared the binding of SP to the NTS_1_R bound to neurotensin 8-13^40^, the μ-opioid receptor bound to the peptide mimetic agonist DAMGO^41^, and the orexin 2 receptor bound to orexin B^42^ (Supplementary Fig. 5). All of these neuropeptides make extensive contacts with the deep 7TM pocket, likely important for determining their efficacy as agonists for their respective receptors. The extended conformations of the peptides in the receptor binding pockets enable further interactions with the extracellular loops. In contrast to the binding of these other neuropeptides at their cognate receptors, SP makes more extensive contacts with ECL2 and the N-terminus of NK1R, manifesting as an outward displacement of the extracellular tip of TM1 and a more ordered N terminus (Supplementary Fig. 5).

The SP orthosteric binding pocket is distinct from the binding sites of NK1R antagonists determined in previous inactive-state structures^38,43,44^. For example, the antagonist netupitant^38^ (PDB: 6HLP) minimally overlaps with SP (Fig. 2B), with only the 2-methylphenyl and 3,5-bis(trifluoromethyl)phenyl groups of netupitant binding in the same region as M11 of SP (Fig. 2B). The core of the netupitant antagonist scaffold, however, extends along TM4 and TM5 toward the extracellular region of the receptor in a portion of the orthosteric pocket that is not occupied by SP. All structurally characterized NK1R antagonists possess a similar molecular scaffold to netupitant and bind to a relatively small portion of the total SP binding site (Supplementary Fig. 5). This distinct binding topology is consistent with prior mutagenesis data, which found only two NK1R residues, Q165^4.60^ and Y287^7.35^, are important for both SP and non-peptide antagonists binding^9,45–49^ (Supplementary Fig. 5).

The expansive SP-NK1R interface is consistent with prior mutagenesis efforts, which found that residues both within the deep 7TM site and the NK1R N-terminus potently reduce SP binding affinity^9,47,49^. Our structure of the SP-NK1R complex revealed that the amidated C-terminus of SP forms an extensive hydrogen-bonding network with NK1R residues N85^2.57^, N89^2.61^, H108^3.28^, and Y287^7.35^ (Fig. 2D). To finely probe the importance of specific hydrogen bonds in SP binding to NK1R, we tested the ability of SP to activate NK1R mutants with conservative amino acid substitutions at these key positions in a Ca^2+^ mobilization assay. In contrast to the dramatic loss of potency previously observed with non-conservative alanine mutations at these sites^49^, we observed relatively minor changes in SP potency or maximal efficacy with these conservative mutations (Fig. 2E and Supplementary Fig. 6).

We hypothesized that other SP-NK1R interactions, perhaps those in the extracellular regions of the receptor, could compensate for the disrupted hydrogen bonding network in the deep portion of the NK1R pocket. To test this, we examined the potency and efficacy of a truncated version of SP containing only residues 6-11 (SP6-11, Fig. 3A), which would be unable to interact with the NK1R ECL2 and N-terminus. As observed previously^18^, we found that SP6-11 is equally potent as SP in stimulating Ca^2+^ signaling and IP1 accumulation at wild-type NK1R (Fig. 3B and Supplementary Fig. 6). When tested against our conservative NK1R mutants targeting the SP C-terminal amide hydrogen bonding network, we observed a dramatic 30-1000 fold loss in potency for SP6-11 (Fig. 2E). We therefore conclude that the extensive hydrogen bonding network recognizing the amidated C-terminus of SP is indeed important for Ca^2+^ signaling, but that the extended contacts between SP and NK1R in the extracellular regions serve an important role in NK1R signaling beyond simply dictating the tachykinin receptor subtype selectivity of SP.

**Figure 3.**
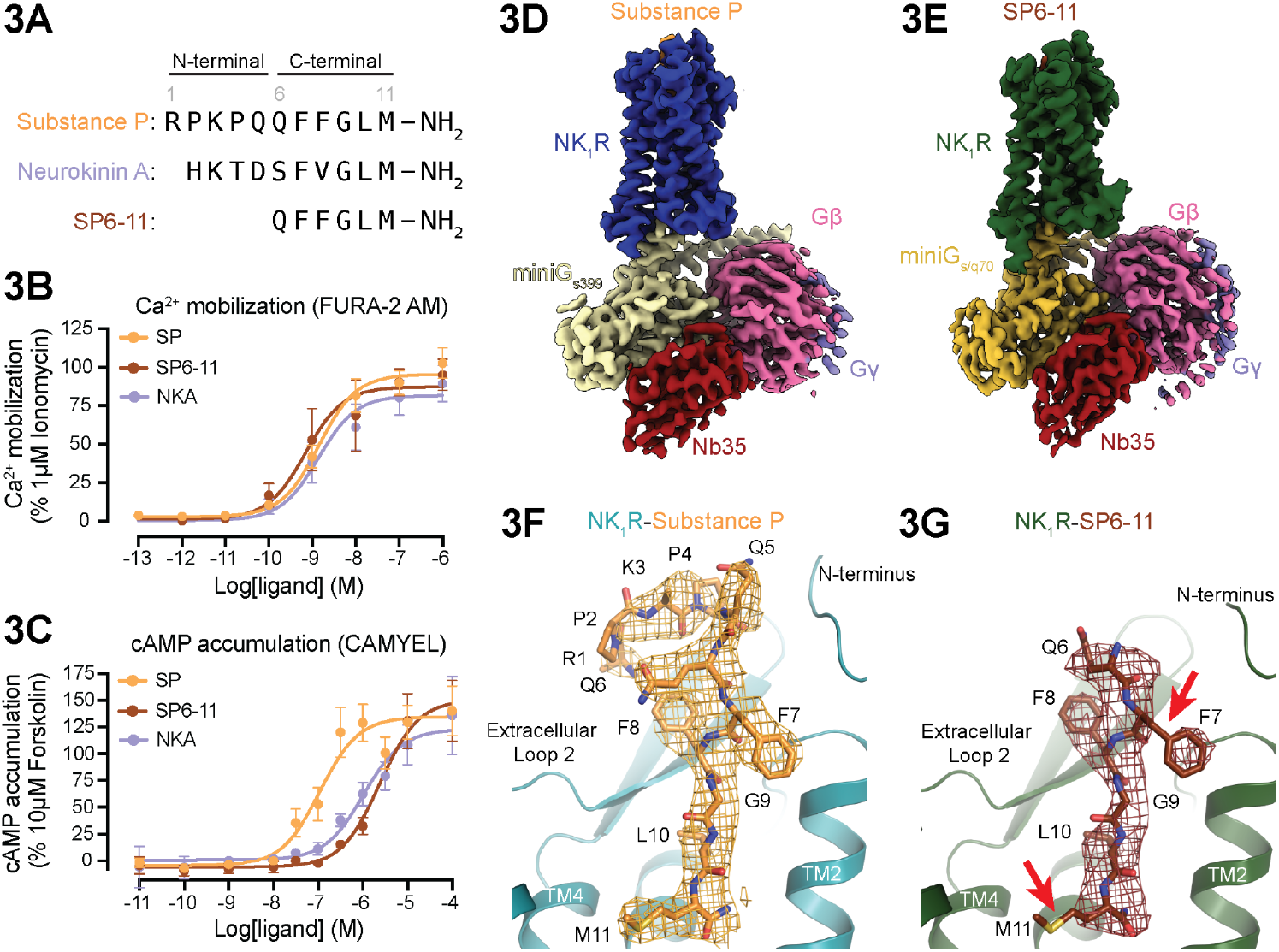
Structural interrogation of SP6-11, a G_q_-selective tachykinin. **(A)** Sequence of SP, Substance P 6-11 (SP6-11) and Neurokinin A (NKA). **(B,C)** Ca^2+^ and cAMP signaling assays demonstrate that SP6-11 and NKA signal potently through G_q_ but have diminished potency for G_s_ signaling. Signaling graphs represent the global fit of grouped data ± s.e.m. from n ≥ 3 independent biological replicates. Full quantitative parameters from this experiment are listed in Supplementary Table 2. **(D)** Cryo-EM map of SP-NK1R-miniG_s399_-Nb35 complex. **(E)** Cryo-EM map of SP6-11-bound NK1R-miniG_s/q70_-Nb35 complex. **(F,G)** Unsharpened density maps at equivalent enclosed volume thresholds for SP and SP6-11 are shown as mesh. For SP6-11, density for the M11 side chain is absent, as is connecting density for the F7 side chain (indicated by red arrows). By contrast, these regions are clearly resolved for SP. Density map for the peptide is contoured 2 Å away from modeled atoms.

### Structural interrogation of NK1R signaling bias

While SP potently activates both G_q_ and G_s_ signaling downstream of NK1R^22^, NKA and N-terminally truncated SP analogs are weaker agonists of G_s_ signaling^17,18^. We confirmed these prior results in signaling studies. SP, NKA, and SP6-11 produced equally potent and efficacious Ca^2+^ and IP signaling responses (Fig. 3B and Supplementary Fig. 6). By contrast, NKA and SP6-11 were 6- and 16-fold less potent than SP in eliciting cAMP accumulation, respectively (Fig. 3C, Supplementary Table 2), confirming their G_q_ selective signaling profiles.

Prior pharmacology studies have demonstrated that SP and its analogs bind to NK1R in two distinct conformations that likely depend on the specific G protein coupled to the receptor^24,50^. Indeed, NK1R-G_q_and NK1R-G_S_ fusion proteins display different binding affinities for SP^50^, suggesting that G_q_- and G_s_-coupled NK1R exist in distinct conformations. We therefore reasoned that additional cryo-EM structures of active NK1R may provide insight into how SP and other tachykinins induce distinct G_q_ and G_s_ signaling outcomes. In particular, we speculated that differences in the SP-ECL2 interaction interface may explain the diminished G_s_ agonism of NKA and SP6-11. To explore how SP induces G_s_ signaling, we determined a structure of SP-NK1R bound to a miniG protein analog of G_s_ (miniG_s399_) at 3.1 Å resolution (Fig. 3D, Supplementary Fig. 1, Supplementary Fig. 7). Furthermore, to understand how NKA and SP6-11 induce G_q_ selective NK1R signaling, we determined the structure of a SP6-11-NK1R-miniG_s/q70_ complex at 3.2 Å resolution (Fig. 3E, Supplementary Fig. 1, Supplementary Fig. 8).

The structure of the full-length SP-NK1 R-miniGs_399_ complex is almost identical to our SP-NK1R-miniG_s/q70_ complex, with overall NK1R root mean square deviation (RMSD) of 0.38 Å. Importantly, we observe clearly resolved EM density for the N-terminus of SP, which interacts with the NK1R extracellular regions in a very similar manner to the SP-NK1 R-miniG_s/q70_ structure (Supplementary Fig. 9). The similarity in these SP-NK1R-miniG protein structures suggests that the overall conformation of G_q_- and G_s_-coupled NK1R may be similar while bound to SP. However, there are important caveats to this interpretation. First, the primary interaction between NK1R and the miniG proteins is the insertion of the G protein C-terminal α5-helix into the receptor 7TM core. In the chimeric miniG_s/q70_ protein, the α5-helix is derived from G_q_, whereas the remainder of the miniG_s/q70_ protein is derived from G_s_. It is possible that the interaction between NK1R and wild-type G_q_ may be different from what we observe with miniG_s/q70_ here. Furthermore, both miniG proteins used here have been engineered to stabilize the active G protein conformation and increase the affinity of the G protein-GPCR interaction. We cannot rule out the possibility that these modifications may influence the specific conformation of NK1 R that we observed in our structures. Finally, our cryoEM reconstructions capture a single, low-energy state of a nucleotide-free NK1R-miniG protein complex. It is also possible that G_q_ and G_s_ signaling selectivity arises from transient NK1R G protein-coupled states that are not structurally observed in our work.

The structure of SP6-11 activated NK1R-miniG_s/q70_ also revealed a highly similar receptor-G protein conformation when compared to full-length SP, with notable exceptions in the peptide binding site (Fig. 3F,G and Supplementary Fig. 9). We observed a shorter density for SP6-11 in the orthosteric binding pocket, consistent with the N-terminal truncation of SP. The cryo-EM density for SP6-11 is comparatively worse than the density for both full-length SP reconstructions when viewed with unsharpened maps at the same enclosed volume threshold (Fig. 3G). Specifically, there is a lack of continuous electron density between the peptide backbone and the F7 sidechain and the electron density for the M11 sidechain is completely missing. By contrast, the density for the receptor is comparatively well resolved for both full-length SP and SP6-11 structures, suggesting that the weaker density we observed for SP6-11 does not arise from local resolution artifacts (Supplementary Fig. 9). Selectively weaker density for F7 and M11 may arise from increased dynamic motion of SP6-11 compared to full-length SP. We speculate that increased SP6-11 dynamic motion may result from a lack of stabilizing contacts between the N-terminus of SP and the extracellular regions of NK1R, potentially leading to both increased sensitivity of SP6-11 to mutations in the deep 7TM pocket (Fig. 2E) and decreased potency of SP6-11-mediated G_s_-signaling (Fig. 3C).

### Truncation of SP N-terminus increases C-terminus mobility

We turned to all-atom molecular dynamics (MD) simulations to understand the mechanism by which SP N-terminal truncation leads to G_q_ selective signaling. To this end, we performed 12 independent 2-μs simulations of active NK1R bound to SP and another 12 of active NK1R bound to SP6-11.

In our simulations, SP6-11 is less restrained in its motion than the corresponding C-terminal residues of SP and explores more space within the binding pocket (Fig. 4). SP6-11 residues F7 and M11 exhibit particularly notable differences in dynamics compared to SP. In simulations of NK1R bound to full-length SP, the F7 side chain remains mostly between TM7 and TM2, as in the SP-bound NK1R structure (Fig. 4A). In simulations with SP6-11, on the other hand, the F7 side chain samples a wider range of orientations (Fig. 4C). The side chain of M11 adopts two major orientations in simulations of SP: one pointing between TM5 and TM6 as in the SP-bound structure of NK1 R and one pointing between TM6 and TM7. For SP6-11, we observe a wider range of M11 side chain orientations with additional conformations pointing towards TM6 and TM7 (Fig. 4C). Altogether, we observe a significant increase in the root mean square fluctuation (RMSF) for SP6-11 bound to NK1R, both for the entire C-terminal peptide region and for the F7 and M11 residues (Fig. 4B). This increased flexibility is consistent with poorly resolved regions for the SP6-11 peptide in our cryo-EM structure (Fig. 3G). We conclude that disruption of the interactions between the SP N-terminus and the NK1R leads to destabilization of SP C-terminal residues.

**Figure 4.**
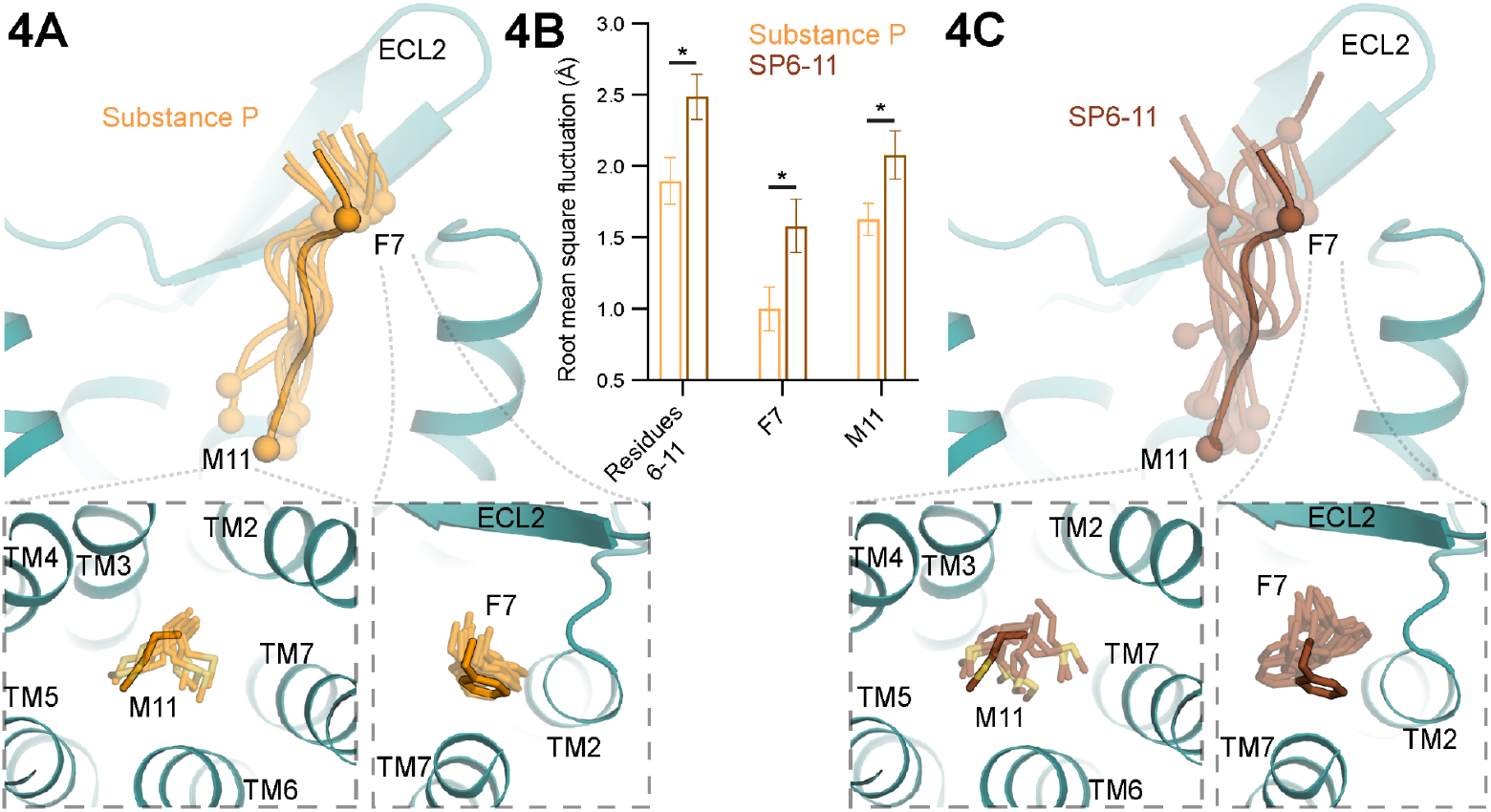
Molecular dynamics shows increased mobility of SP6-11. **(A)** Molecular dynamics simulation snapshots for backbone of C-terminal residues of Substance P (SP). Cα atoms for methionine 11 (M11) and phenylalanine 7 (F7) are shown as spheres. The starting cryo-EM structure of SP in the binding pocket is shown with outline; simulation snapshots are transparent. Simulations included all 11 amino-acids of SP, but only residues 6-11 are shown here to enable comparison with SP6-11. Insets show conformations for F7 and M11 side chains of SP. **(B)** Quantitation of peptide mobility in molecular dynamics simulations as measured by root mean square fluctuation (RMSF). Bar graphs show mean RMSF ± s.e.m. from twelve independent molecular dynamics simulations (* = p < 0.05, two-sided Welch’s t-test). **(C)** Simulation snapshots for SP6-11. Insets show alternative conformations for F7 and M11 side chains of SP6-11.

Intriguingly, the different orientations observed for M11 and F7 affect contacts with TM7. Given that active NK1R is already in an unusual ‘non-canonical’ conformation, with TM7 inactive but TM6 in an outward position, it is tempting to speculate that these different interactions with TM7 could also stabilize distinct intracellular conformations differing in TM7 conformation. Different TM7 conformations have been previously shown to mediate differential signaling responses of another peptidergic GPCR, the angiotensin II type 1 receptor^51^.

### Disruption of SP contacts with ECL2 biases signaling

Our simulations suggested that the interactions of the N-terminal region of SP with NK1 R ECL2 serve to stably position the C-terminus of the peptide. Disruption of these interactions may therefore destabilize the C-terminal region of SP and achieve similar G_q_ preferential signaling as observed for SP6-11. To directly test the relevance of interactions between ECL2 and SP, we designed NK1R mutations that disrupt the SP-ECL2 interface (Supplementary Fig. 10). Two such mutations, M174I and R177M, displayed G_q_ preferential signaling by SP. In structures of NK1R bound to SP, M174 makes hydrophobic contacts with F8 of SP while R177 forms an extended hydrogen-bond network with the SP backbone and NK1R residues N96^2.68^ in TM2 and N23 in the N-terminus (Fig. 5A). In signaling studies, both M174I and R177M are equally potent and efficacious as wild-type NK1R at Ca^2+^ mobilization (Fig. 5B). By contrast, both of these mutations significantly decrease cAMP production by SP, with R177M displaying a 20-fold reduction in potency and a >3-fold reduction in efficacy compared to wild-type NK1R (Fig. 5C, Supplementary Table 2). Importantly, these mutants are expressed at similar levels as wild-type NK1R (Supplementary Fig. 10). Contacts between SP and NK1R ECL2 are therefore critically important for potent G_s_-coupled cAMP signaling.

**Figure 5.**
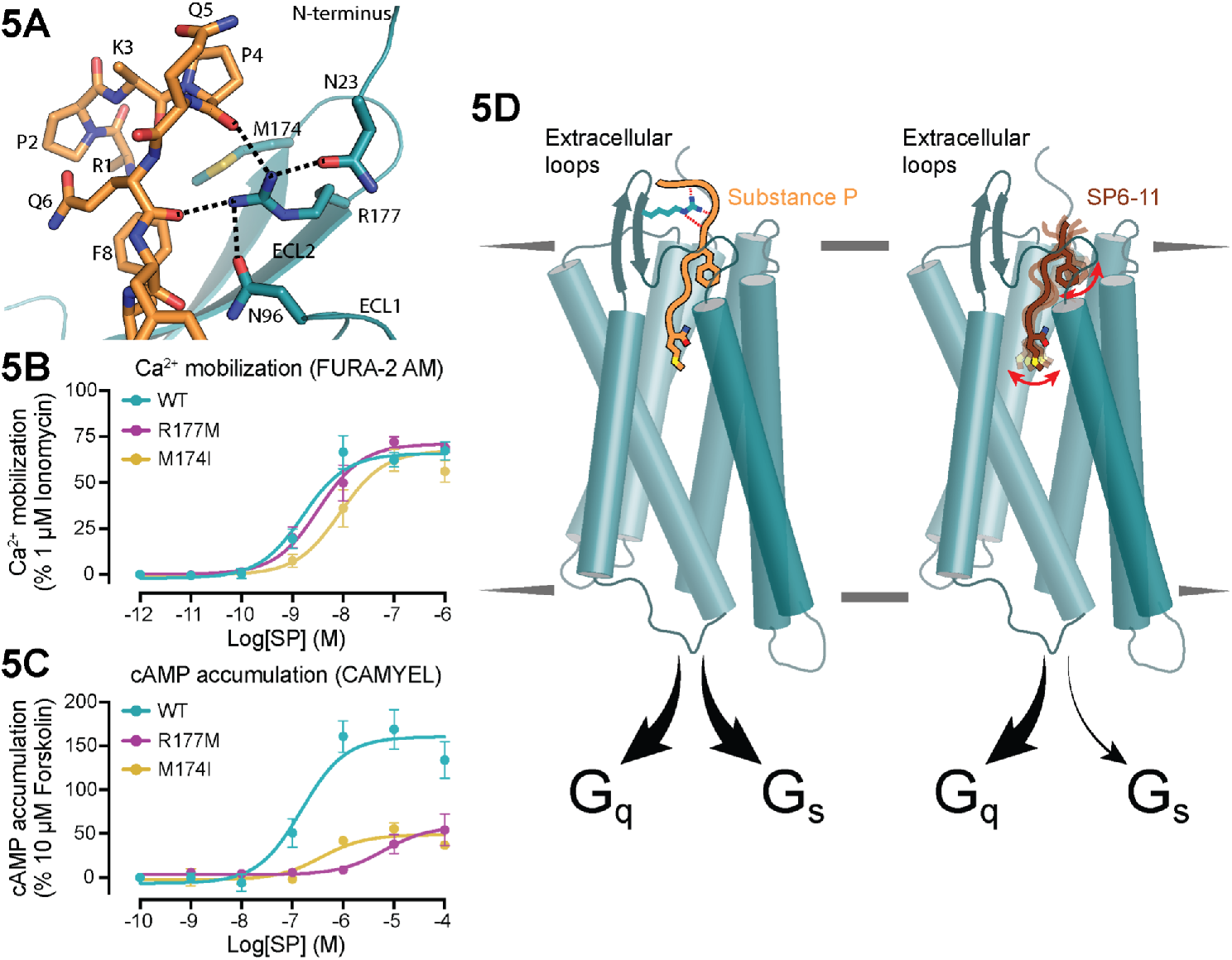
Disruption of SP-NK1R ECL2 contacts leads to G_q_-selective signaling. **(A)** NK1R ECL2 contacts with the N-terminal region of SP. R177 engages in an extended hydrogen bonding network with the SP backbone while M174 makes van der Waals contacts with R1 and P4 **(B, C)** Ca^2+^and cAMP signaling assays for point mutants disrupting ECL2-SP interactions. Disruption of SP-NK1R ECL2 contacts leads to strongly G_q_ selective signaling. Signaling graphs represent the global fit of grouped data ± s.e.m. from n > 3 independent biological replicates. Full quantitative parameters from this experiment are listed in Supplementary Table 3. **(C)** Model for tuning of G protein selectivity driven by contacts between SP and ECL2.

## Conclusion

Substance P is a prototypical member of the broader family of neuropeptides that act at GPCRs to modulate neuronal function. Our structures of full-length SP bound to active NK1R revealed an extensive contact interface with NK1R, stretching from the deeply buried regions in the 7TM domain to the extracellular regions of the receptor. A network of specific hydrogen bonds between the amidated C-terminus of SP and the deep orthosteric pocket of NK1R are important for peptide recognition; removal of specific hydrogen bonds impairs the ability of SP6-11 to activate NK1R. Our structures also reveal important contacts between the SP N-terminus and the extracellular loops of NK1R, providing insight into how less-conserved sequences in neuropeptides engage GPCRs. While our structural views of full-length neuropeptides bound to their cognate GPCRs remains limited to only a few other examples^42,52^, an emerging theme is that peptides bind in an extended manner, with regions of the peptide engaging the receptor extracellular loops.

Our work revises the ‘message-address’ model for peptidergic signaling at GPCRs. In particular, we demonstrate that interactions between the N-terminal region of SP and the extracellular loops of NK1R, which were previously characterized as conferring receptor subtype selectivity, are also required for balanced signaling via both the G_q_ and G_s_ signaling pathways. Loss of these interactions, due to either truncation of SP’s N-terminus or NK1R ECL2 mutations, leads to G_q_-selective signaling. Other endogenous tachykinins that signal selectively via G_q_, such as NKA, likely do so because they lack sequences that can engage the NK1R extracellular loops. Neuropeptide regions that engage GPCR extracellular loops may therefore specify not only which receptor subtype a peptide preferentially engages but also the signaling outcomes downstream of a specific GPCR.

Our work has broader implications for other neuropeptide GPCRs. Pharmacological studies with other neuropeptides, including opioid peptides^53,54^, neuropeptide S^55,56^, and neuropeptide Y^57^ suggest that interactions between the divergent, less conserved regions of the neuropeptide and their cognate receptor extracellular loops, which also diverge in sequence, can tune signaling efficacy via multiple G proteins or β-arrestins. Our work highlights the role of peptide-receptor extracellular contacts in determining the conformational flexibility of the core “message” region of SP. A similar mechanism may be responsible for fine-tuning intracellular signaling, or for promoting a complete signaling response when multiple endogenous neuropeptides are present and acting at a single GPCR. Assessing the structure and dynamics of neuropeptides bound to their cognate receptors thus promises to yield a mechanistic understanding of what drives GPCR signaling complexity, and may eventually provide a path to control such complex signaling with designed molecules.

## METHODS

### Expression and purification of full-length Substance P-NK1R-miniG_s/q70_ complex

For structure determination, human *TACR1* with an N-terminal HA signal sequence followed by a FLAG epitope tag was cloned into a pcDNA^™^3.1/Zeo^(+)^ vector containing a tetracycline-inducible expression cassette. The miniG_s/q70_ protein^25^ was fused to the NK1R C-terminus, preceded by a flexible glycine/serine linker and rhinovirus 3C protease recognition site (LEVLFQGP). The resulting NK1R-miniG_s/q70_ fusion construct was transfected into adherent Expi293F™ Inducible Human Embryonic Kidney Cells (unauthenticated and untested for mycoplasma contamination, Life Technologies) using Lipofectamine 2000^Ω^ and cells were maintained in DMEM (Gibco, 11995-065) + 10% FBS (Gibco), 100 U/mL penicillin and 100 μg/mL streptomycin at 37 °C and 5% CO_2_ in a standing incubator. Cells stably incorporating the NK1R-miniG_s/q70_ fusion plasmid were selected under antibiotic pressure with zeocin (500 μg/mL) and blasticidin (10 μg/mL). The resulting polyclonal Expi293F™ NK1R-miniG_s/q70_ stable cell line was then adapted to suspension culture and maintained in Expi293™ Expression Medium (Gibco) supplemented with zeocin (5 μg/mL) and blasticidin (5 μg/mL) at 37 °C and 8 % CO_2_ on a shaking platform at 125 rpm. Expression of NK1R-miniG_s/q70_fusion protein was induced with addition of 4 μg/mL doxycycline hyclate (Sigma Aldrich) and enhanced with 20 mM sodium butyrate (Sigma Aldrich). Two liters of induced Expi293F™ NK1R-miniG_s/q70_stable cells were harvested 24 hours after induction and stored at −80 °C until further use.

For purification, cells were thawed and washed with hypotonic buffer (20 mM HEPES pH 7.5, 1 mM EDTA) supplemented with protease inhibitors (20 μg/mL leupeptin, 160 μg/mL benzamidine, 1 mM PMSF), reducing agent (100 μM TCEP) and 100 nM Substance P (Tocris). The membrane fraction was then solubilized with 50 mM HEPES pH 7.5, 300 mM NaCI, 1% (w/v) lauryl maltose neopentyl glycol (L-MNG, Anatrace), 0.1% cholesteryl hemisuccinate (CHS, Steraloids), protease inhibitors, 100 μM TCEP, 5 mM ATP, 2 mM MgCl_2_, and 1 μM Substance P for 1.5 hours at 4 °C. After high-speed centrifugation, the supernatant was subjected to affinity purification using homemade M1 anti-FLAG antibody coupled to Sepharose beads. NK1R-miniG_s/q70_ bound to M1-beads was washed extensively to gradually decrease detergent and salt concentration and was eluted in 50 mM HEPES pH 7.5, 150 mM NaCI, 0.0075% (w/v) L-MNG, 0.0025% (w/v) glyco-diosgenin (GDN, Anatrace), 0.001% CHS, 100 μM TCEP, 100 nM Substance P, 5 mM EDTA, and 0.2 mg/mL FLAG peptide (Genscript). Eluted NK1R-miniG_s/q70_ was concentrated with a 50 kDa MWCO spin concentrator (Millipore) and purified to homogeneity with size-exclusion chromatography, using a Superdex S200 Increase 10/300 GL column (GE Healthcare) equilibrated in 20 mM HEPES pH 7.5, 150 mM NaCI, 0.0075% (w/v) L-MNG, 0.0025% (w/v) GDN, 0.001% CHS, 100 μM TCEP, and 100 nM Substance P. Fractions containing monodisperse NK1R-miniG_s/q70_ fusion protein were pooled, mixed with 2.5x molar excess of Gβ_1_γ_2_ heterodimer, Nb35, and Substance P, and incubated overnight at 4 °C. The next day, the heterotrimeric complex was concentrated with a 50 kDa MWCO spin concentrator and excess Gβ_1_γ_2_ and Nb35 was removed via size-exclusion chromatography, using a Superdex S200 Increase 10/300 GL column (GE Healthcare) equilibrated in 20 mM HEPES pH 7.5, 150 mM NaCI, 0.00075% (w/v) L-MNG, 0.00025% (w/v) GDN, 0.0001% CHS, 100 μM TCEP, and 100 nM Substance P. Resulting SP-NK1R-miniG_s/q70_ heterotrimeric complex was concentrated with a 50 kDa MWCO spin concentrator to 1.93 mg/mL (14 μM) for preparation of cryo electron microscopy grids.

### Expression and purification of Substance P (6-11)-NK1R-miniG_s/q70_ complex

The SP6-11-bound NK1R-miniG_s/q70_ fusion protein was expressed and purified exactly as described above for the full-length SP-NK1R-miniG_s/q70_ fusion protein, with the exception of replacing all SP incubations with SP6-11 throughout the purification. Incubation of SP6-11-NK1R-miniG_s/q70_ fusion protein with Gβ_1_γ_2_ and Nb35 was performed as described above for the full-length SP complex. The resulting SP6-11-NK1R-miniG_s/q70_ heterotrimeric complex was concentrated with a 50 kDa MWCO spin concentrator to 2.91 mg/mL (21 μM) for preparation of cryo electron microscopy grids.

### Expression and purification of full-length Substance P-NK1R-miniG_s399_ complex

The NK1R-miniG_s/q70_ fusion construct (generation described above) was modified to replace the miniG_s/q70_ protein with the miniG_s399_^33,34^ protein using Gibson cloning. The subsequent NK1R-miniG_s399_fusion construct was transiently transfected into 200-mLs of Expi293F™ Inducible Human Embryonic Kidney Cells (unauthenticated and untested for mycoplasma contamination, Life Technologies) using the Expifectamine Transfection Kit (Life Technologies), following the manufacturer’s instructions. Expression of the NK1R-miniG_s399_fusion protein was induced and enhanced 18 hours after transfection with addition of 1 μg/mL doxycycline hyclate (Sigma Aldrich), 10 mM sodium butyrate (Sigma Aldrich), and addition of enhancers from the Expifectamine Transfection Kit, as per the manufacturer’s instructions. Cells were harvested 24 hours after induction and stored at −80 °C until further use.

The full-length SP-NK1R-miniG_s399_ fusion protein was purified exactly as described above for the full-length SP-NK1R-miniG_s/q70_ fusion protein. Incubation of SP-NK1R-miniG_s399_ fusion protein with Gβ_1_γ_2_ and Nb35 was performed exactly as described above for the SP-NK1 R-miniG_s/q70_ complex. The resulting SP-NK1R-miniG_s399_ heterotrimeric complex was concentrated with a 50 kDa MWCO spin concentrator to 2.86 mg/mL (20 μM) for preparation of cryo electron microscopy grids.

### Expression and purification of Gβ_1_γ_2_

The Gβ_1_γ_2_ heterodimer was expressed in *Tnchoplusia ni* (Hi5) insect cells (Expression Systems, unauthenticated and untested for mycoplasma contamination) using a single baculovirus, as previously described^60^. A single bicistronic baculovirus encoding the human Gβ_1_ subunit with a N-terminal 6x His-tag and rhinovirus 3C protease site and untagged human Gγ_2_ subunit was generated using the BestBac method (Expression systems) in *Spodoptera frugiperda* (Sf9) insect cells (Expression Systems 94-001F, unauthenticated and untested for mycoplasma contamination). Hi5 insect cells were transduced with baculovirus at a density of ~3.0 x 10^6^ cells/mL, grown at 27 °C and shaking at 130 rpm. Cultures were harvested 48 hours after transduction, and cell pellets were stored at −80 °C until further use. Frozen cell pellets were thawed and washed in a hypotonic buffer containing 10 mM Tris pH 8.0, 5 mM β-mercaptoethanol (β-ME), and protease inhibitors (20 μg/mL leupeptin, 160 μg/mL benzamidine). The membrane fraction was collected by centrifugation and then solubilized with 20 mM HEPES pH 7.5, 100 mM NaCI, 1% (w/v) sodium cholate, 0.05% dodecyl maltoside (DDM, Anatrace), 0.005% cholesteryl hemisuccinate (CHS, Steraloids), 5 mM β-ME, protease inhibitors, and 5 mM Imidazole for 1 hour at 4 °C. After high-speed centrifugation, the supernatant was subjected to affinity purification with HisPur^™^ Ni-NTA resin (Thermo Scientific). Bound Gβ_1_γ_2_ heterodimer was washed extensively and detergent was slowly exchanged to 0.1 % (w/v) lauryl maltose neopentyl glycol (L-MNG, Anatrace) and 0.01% CHS before elution with 20 mM HEPES pH 7.5, 100 mM NaCI, 0.1% L-MNG, 0.01% CHS, 270 mM imidazole, 1 mM dithiothreitol (DTT), and protease inhibitors. Eluted Gβ_1_γ_2_ heterodimer was pooled and 3C protease was added to cleave the N-terminal 6x His-tag. The resulting Gβ_1_γ_2_ heterodimer was dialyzed overnight in 20 mM HEPES pH 7.5, 100 mM NaCI, 0.02% L-MNG, 0.002% CHS, 1 mM DTT, and 10 mM imidazole. Reverse Ni-NTA affinity chromatography was performed to remove uncleaved heterodimer. The resulting Gβ_1_γ_2_ was then incubated for 1 hour at 4 °C with lambda phosphatase (New England Biolabs), calf intestinal phosphatase (New England Biolabs), and antarctic phosphatase (New England Biolabs) to dephosphorylate the protein. Gβ_1_γ_2_ was further purified by anion exchange chromatography using a MonoQ4.6/100 PE (GE Healthcare) column. The resulting protein was pooled and dialyzed overnight in 20 mM HEPES pH 7.5, 100 mM NaCI, 0.02% L-MNG, and 100 μM TCEP, concentrated with a 3 kDa centrifugal concentrator. Glycerol was added to a final concentration of 20%, and the protein was flash frozen in liquid N_2_and stored at −80 °C until further use.

### Expression and purification of Nb35

Nanobody35 (Nb35)^26^ with a N-terminal pelB signal sequence and a C-terminal Protein C affinity tag (EDQVDPRLIDGK) was cloned into a pET-26b IPTG-inducible bacterial expression vector. This vector was transformed into BL21 Rosetta *Escherichia coli* cells and grown overnight in Luria Broth supplemented with 50 μg/mL kanamycin shaking at 225 rpm and 37 °C. Next day, the saturated overnight culture was used to inoculate 8 L of Terrific Broth (supplemented with 0.1% glucose, 2 mM MgCl_2_, and 50 μg/mL kanamycin) and cells were grown shaking at 225 rpm at 37 °C. When cells reached an OD_600_ = 0.6, expression of Nb35 was induced with addition of 400 μM IPTG and the temperature was reduced to 20 °C for 21 hours. Cells were harvested by centrifugation and stored in the −80 °C until further use. For purification of Nb35, cells were thawed and resuspended in SET Buffer (200 mM Tris pH 8.0, 500 mM sucrose, 0.5 mM EDTA) supplemented with protease inhibitors (20 μg/mL leupeptin, 160 μg/mL benzamidine) and benzonase. After 30 minutes of stirring, two equal volumes of miliQ H_2_O were added to initiate hypotonic lysis. After 45 minutes of stirring, NaCI was added to 150 mM, CaCl_2_ was added to 2 mM, and MgCl_2_ was added to 2mM. Insoluble matter was then separated by high speed centrifugation and the supernatant was subjected to affinity purification with homemade anti-Protein C antibody coupled to Sepharose beads. After extensive washing, bound Nb35 was eluted with 20 mM HEPES pH 7.5, 100 mM NaCI, and 2 mM CaCl_2_, 0.2 mg/mL Protein C peptide, and 5 mM EDTA pH 8.0. Eluted Nb35 was collected, concentrated, and injected over an Superdex S75 Increase 10/300 GL column (GE Healthcare) size-exclusion chromatography column equilibrated in 20 mM HEPES pH 7.5, 100 mM NaCI. Fractions containing Nb35 were collected, concentrated, glycerol was added to a final concentration of 20%, and aliquots of Nb35 were flash frozen in liquid nitrogen and stored in the −80 °C until further use.

### Cryo-EM sample vitrification and data collection

For cryogenic electron microscopy, 3 μL of the SP-NK1R-miniG_s/q70_ heterotrimeric complex at 13.8 μM was added to 300 Mesh 1.2/1.3R Au Quantifoil grids previously glow discharged at 15 mA for 30 seconds with a Pelco easiGlow Glow discharge cleaning system. Grids were blotted with Whatman No. 1 qualitative filter paper in a Vitrobot Mark IV (Thermo Fisher) at 8°C and 100% humidity for 1 second using a blot force of 4 prior to plunging into liquid ethane. The SP-NK1R-miniG_s399_ and SP6-11-NK1R-miniG_s/q70_ heterotrimeric complexes were frozen under identical blotting conditions at 20 μM and 20.7 μM, respectively.

For the SP-NK1R-miniG_s/q70_ heterotrimeric complex, 3,755 super-resolution movies were recorded with a 300 keV Titan Krios (Thermo Fisher) equipped with a K3 detector and BioQuantum energy filter (Gatan) with a zero-loss energy selection slit width set to 20 eV and a defocus range of −0.8 to −2.0 μm. Each dose-fractionated 120-frame movie was collected at a dose rate of 8.0 e^-^pix^-1^s^-1^ for 5.9 seconds at a nominal magnification of 105,000x (physical pixel size of 0.835 Å pix^-1^) resulting in a cumulative dose of 67 e^-^ Å^-2^. Exposure areas were acquired with automated scripts in a 3×3 image shift collection strategy using SerialEM.^61^ The 3878 super-resolution movies of the SP6-11-NK1R-miniG_s/q70_ heterotrimeric complex were acquired with identical acquisition settings on the same 300 keV Titan Krios as previously described.

For the SP-NK1R-miniG_s399_ heterotrimeric complex, 3670 dose-fractionated movies were collected in counting mode with a 300 keV Titan Krios equipped with a K3 detector and BioQuantum energy filter (Gatan) with a zero-loss energy selection slit width set to 20 eV and a defocus range of −0.8 to −2.1 μm. At a dose rate of 24 e^-^pix^-1^s^-1^, each 1.52s exposure was fractionated across 60 frames for a total dose of 49.4 e^-^ Å^-2^. Data were collected using aberration free image shift (AFIS) with EPU 2.10.

### Cryo-EM Image Processing

During data collection, movies of the NK1R-miniG_s/q70_ complex were motion corrected and dose weighted with UCSF MotionCor2^62^ and binned to physical pixel size. Post-acquisition, micrographs were imported into cryoSPARC^63^ for contrast transfer function determination via patch CTF. 7,329,811 particles were template picked with 20 Å low-pass filtered projections of the NK1 R-miniG_s/q70_ heterotrimeric complex from a prior screening collection on a 200 keV Talos Arctica. 3,555 micrographs comprising 6,945,760 particles were curated for further processing via CTF fit estimated resolution, ice thickness, and particle pick power scores. Particles were extracted in a 72-pixel box, Fourier cropped from 288 pixels. 200 2D class averages were generated using a maximum alignment resolution of 8 Å, 30 online-EM iterations, and 200 particles per class during each online EM iteration. 1,901,254 particles were selected from qualitatively “good” classes containing any averages that showed “multi-lobed” densities or appeared to be “top” or “bottom” views. Following 2D classification, 500,000 selected particles were used for *ab initio* reconstruction into 3 classes. A single “multi-lobed” class suggestive of an intact NK1R-miniG_s/q70_ heterotrimeric complex was selected and used alongside three poorly aligned “junk” classes generated from *ab initio* volumes from <100 particles. All particles selected from 2D classification as described above were subject to 3D classification with alignment against the NK1R-miniG_s/q70_ and poorly aligned classes from the earlier *ab initio* runs. 718,386 particles in the NK1R complex class were re-extracted without Fourier cropping and subject to the same 3D classification scheme with a new *ab initio* NK1R-miniG_s/q70_ heterotrimeric complex class generated from 300,000 particles and three “junk” classes. Using UCSF pyEM^64^, 589,973 particles in the NK1R-miniG_s/q70_ class were exported from cryoSPARC for alignment-free classification in RELION^65^ into 4 classes for 50 iterations with a *τ* parameter of 8. Two classes containing 122,222 particles were imported into cisTEM^66^ for manual “focused” refinements utilizing whole complex and upper-transmembrane domain masks, respectively. To generate local resolution estimates, the NK1R-miniG_s/q70_ complex mask and unfiltered half maps were imported into cryoSPARC. The same half maps and mask were used for directional FSC curve generation^67^.

3,670 SP-NK1R-miniG_s399_ movies were motion-corrected post acquisition with UCSF MotionCor2. CTF estimation and template-based automatic particle picking were performed in cryoSPARC. 4,865,341 particles were picked using templates generated from 2D class averages of SP-NK1R-miniG_s399_ as determined from an earlier screening collection. 4,256,322 particles were extracted in an 80 pixel box after curating micrographs via CTF fit estimated resolution, ice thickness, and particle pick power scores. Extracted particles were subject to a round of 3D classification with alignment (Heterogeneous refinement) using a 20Å low-pass filtered initial volume of NK1R-miniG_s399_ and three additional naïve classes generated from a deliberately under-sampled *Ab initio* job. Particles classified into the NK1R-miniG_s399_ class were selected for further workup. This process was repeated over two additional rounds, decreasing particle Fourier cropping at each subsequent extraction round. 561,901 unbinned particles particles classified into the NK1R-miniG_s399_ complex class were then exported to RELION for alignment-free classification on only 7TM domain features for 25 iterations, 4 classes, *τ* = 8. 288,659 particles in three classes were imported into cisTEM for manual “focused” refinements as in the NK1R-miniG_s/q70_ processing.

3,878 motion-corrected, dose-weighted SP6-11-NK1R-miniG_s/q70_ complex sums were imported into cryoSPARC for CTF estimation and template-based autopicking as previously described. 4,135,538 particles were picked with templates generated from 2D classes determined from a prior screening dataset. Particles were extracted in a 72-pixel box and underwent iterative rounds of 3D classification with alignment on successively unbinned particles classified into a SP6-11-NK1R-miniG_s/q70_ complex class. 553,506 particles were exported to RELION for alignment-free classification on the 7TM domain using the same parameters as previously described, though with six classes instead of four. Visual inspection of each class led to selection of a single class generated from 59,926 particles for manual “focused” refinements in cisTEM using the same masking scheme as for the previously described complexes. Half maps and masks were re-imported into cryoSPARC for GS-FSC determination. dFSC curve calculation utilized the same half maps and masks from focused refinement.

### Model building and refinement: SP-NK1R-miniG_s/q70_

The initial model for NK1R was taken from the high-resolution (2.2 Å) inactive-state structure of NK1R bound to the clinically approved antagonist, netupitant (PDB: 6HLP)^38^. The model was docked into the 3.0 Å EM density map using manual adjustment and the ‘fit in map’ function in UCSF ChimeraX^68^. The initial model was manually rebuilt in Coot and refined with both iterative adjustment in Coot and multiple rounds of global minimization and real space refinement using the Phenix.real_space_refine tool in Phenix^69,70^. In areas of weak sidechain density, residues were capped at the Cβ position to retain sequence information; in areas of weak mainchain density, residues were truncated from the final model. This process was repeated to model the miniG_s/q70_ subunit (starting model: miniG_s_subunit, PDB: 6LI3^30^), Gβ_1_γ_2_ (starting model: Gβ_1_γ_2_ heterodimer, PDB: 3SN6^26^), and Nb35 (starting model: Nb35, PDB: 3SN6^26^). Models were combined into one PDB file in Coot and the model geometry was assessed using Molprobity^71^. Further validation was performed with EMRinger^72^ to compare the map to the model. Finally, map-to-model FSCs were calculated in Phenix^73^. All structure figures were prepared with PyMOL or UCSF ChimeraX^68^.

To model the Substance P peptide, all peptide residues were manually built into the Substance P EM density in Coot. The resulting Substance P model was refined with iterative adjustment in Coot and multiple rounds of global minimization and real space refinement using the real_space_refine tool in Phenix. Due to weak side chain density, R1 and K3 of the peptide are capped at the Cβ position of the residue. To build the amidated C-terminus of M11, a peptide bond connecting the carbonyl-carbon of M11 to a new nitrogen atom was created in Coot. Bond-lengths, bond-angles, dihedral angles, and planes for the new amide moiety were manually adjusted to reflect the accepted values for amide moieties in protein structures.

### Model building and refinement: SP-NK1R-miniG_s399_ & SP6-11-NK1R-miniGs/q70 complexes

The SP-NK1R-miniG_s/q70_ complex model was docked into the 3.1 Å and 3.2 Å EM density maps for the SP-NK1R-miniG_s399_ and SP6-11-NK1R-miniGs/q70, respectively. The models were built and refined as described above. Models were combined into one PDB file in Coot and the model geometry was assessed using Molprobity^71^. Further validation was performed with EMRinger^72^ to compare the map to the model. Finally, map-to-model FSCs were calculated in Phenix^73^. All structure figures were prepared with PyMOL or UCSF ChimeraX^68^.

### Generation of Stable Cell lines and Transfection for signaling assays

Flp-In-HEK293 cells were grown in DMEM supplemented with 5% fetal bovine serum and maintained at 37 °C in a humidified incubator containing 5% CO2. Flp-In-HEK cells were transfected with the pOG44 vector encoding Flp recombinase and the pcDNA5 vector encoding the NK_1_R at a ratio of 9:1 using lipofectamine as the transfection reagent. Twenty-four hours after transfection, the cells were subcultured and forty-eight hours later, the medium was supplemented with 200 μg/ml Hygromycin B as selection agent, to obtain cells stably expressing the NK_1_R.

### Intracellular Ca^2+^ mobilization signaling assay

Flp-In-HEK293 cells stably expressing human NK_1_R WT or mutants were plated in Poly-D-Lysine coated 96-well plates. Cells were washed with calcium buffer (10 mM HEPES, 150 mM NaCI, 2.2 mM CaCl_2_, 1.18 mM MgCl_2_, 2.6 mM KCl, 10 mM D-glucose, 0.5% w/v BSA, 4 mM probenecid, 0.05% v/v pluronic acid F127; pH 7.4) and then loaded with 1 μM Fura-2 AM ester (Life Technologies) in calcium buffer for 45 min at 37°C. Calcium mobilization was measured using a FlexStation 3 plate reader (Molecular Devices). Fluorescence (excitation: 340 nm and 380 nm; emission: 520 nm) was measured at 4 s intervals for 5 cycles. After establishing baseline fluorescence, cells were stimulated with increasing concentrations of the agonists or 1 μM ionomycin (positive control for normalization, to obtain a receptor-independent response) and the response was measured for 17 cycles in SoftMax Pro (v5.4.4) software. GraphPad Prism software (v. 9.0) was used to calculate the area under the curve from the kinetic data and for normalization to vehicle and positive control.

### cAMP accumulation signaling assay

Flp-In-HEK293 cells stably expressing the human NK_1_R WT or mutants were seeded at a density of 2,000,000 cells per 10-cm dish and were transfected the following day using polyethylenimine as the transfection reagent. The cells were transfected with 5 μg CAMYEL biosensor (cAMP sensor using YFP-Epac-RLuc), to allow the detection of cAMP levels by Bioluminescence Resonance Energy Transfer. Twenty-four hours after transfection, the cells were plated into Poly-D-Lysine coated 96-well CulturPlates (PerkinElmer) and grown overnight. The cells were equilibrated in Hank’s balanced salt solution at 37 °C before starting the experiment. Coelenterazine (Promega) was added at a final concentration of 5 μM at least 3 min before measurement. After establishing a baseline response, cells were stimulated with increasing concentrations of the agonists or 10 μM forskolin (positive control for normalization, to obtain a receptor-independent response) and the response was measured for a total of 30 min. The signals were detected at 445-505 nm and 505-565 nm using a LUMIstar Omega instrument (BMG LabTech). GraphPad Prism software (v. 9.0) was used to calculate the area under the curve from the kinetic data and for normalization to vehicle and positive control.

### IP1 accumulation signaling assay

Flp-In-HEK293 cells stably expressing the human NK_1_R WT or mutants were plated in Poly-D-Lysine coated 96-well plates overnight. Cells were equilibrated in Cisbio Bioassays, IP-One Gq kit stimulation buffer (10 mM HEPES, 146 mM NaCI, 4.2 mM KCl, 0.5 mM MgCl_2_, 1 mM CaCl_2_, 5.5 mM D-Glucose, 50 mM LiCI, pH 7.4) for 1 hour prior to agonist stimulation for 1 hour at 37 °C. Cells were then lysed in 25 μl of lysis buffer (50 mM HEPES, 15 mM KF, 1.5% (v/v) Triton-X-100, 3% (v/v) foetal bovine serum, 0.2% (w/v) bovine serum albumin, pH 7.0) and 14 μl of lysis were added to wells of a 384 well white proxiplate (PerkinElmer) for analysis. The Cisbio Bioassays, IP-One competitive immunoassay kit was used to measure myo-Inositol 1 phosphate (IP1) accumulation in cells, based on HTRF^®^ fluorescence resonance energy transfer (FRET) between d2-labeled IP1 (acceptor) and anti-IP1-Cryptate (donor) antibody. These reagents were diluted 1:20 in the lysis buffer and 3 μl of each was added to each well containing the lysates. Lysates were incubated for 1 hour at room temperature before FRET was detected using an Envision plate reader (PerkinElmer). Emission of Lumi4TM-Tb cryptate was detected at 620 nm and emission of d2-conjugated IP1 at 665 nm. Results were calculated from the 665 nm / 620 nm ratio, since the specific signal is inversely proportional to the concentration of IP1, data were transformed and normalised to 1 μM SP in GraphPad Prism software (v. 9.0).

### Enzyme-linked immunosorbent assay (ELISA)

Flp-In-HEK293 cells stably expressing human Flag-NK1R WT, mutants, or untransfected as a control were plated into poly-D-lysine-coated 48-well plates and allowed to adhere overnight. Cells were fixed with 3.7% (v/v) paraformaldehyde in tris-buffered saline (TBS) for 30 min. For total expression, cells were permeabilized by 30-min incubation with 0.5% (v/v) NP-40 in TBS. Cells were then incubated in blocking buffer [1% (w/v) skim milk powder in 0.1 M NaHCO_3_] for 4 hours at room temperature and incubated with mouse M2 anti-FLAG antibody (1:2000, overnight at 4°C). After washing three times with TBS, cells were incubated with anti-mouse horseradish peroxidase-conjugated antibody (1:2000) for 2 hours at room temperature. Cells were washed and stained using the SIGMAFAST OPD substrate (Sigma-Aldrich). Absorbance at 490 nm was measured using an EnVision Multilabel Reader (PerkinElmer). Data were normalized to intact HEK293 cells transfected with NK1R WT.

### Quantification and Statistical Analysis

GraphPad Prism software (v. 9.0) was used for signaling data and statistical analysis. Data points are presented as mean ± standard error of the mean (s.e.m.) based on at least 3 biologically independent experiments with the precise number indicated in Supplementary Table 2 and 3. Data points were normalized to vehicle as 0% and positive control (1 μM ionomycin or 1 μM forskolin) as 100% unless otherwise stated. Concentration-response curves were fitted using the three parameter log(agonist) vs. response equation. pEC50 values were extracted from the curve fit of each individual experiment. Emax was calculated by subtracting Bottom from Top from the curve fit of each individual experiment. Statistical analysis of pEC50 and Emax values was performed using one-way analysis of variance (ANOVA) with Dunnett’s multiple comparison-corrected post hoc test against NK1R WT SP or NK1R WT SP6-11 unless otherwise stated.

### Molecular Dynamics: System setup

We performed simulations of NK1R bound to full-length substance P (SP) and a truncated version of SP reduced to the C-terminal residues 6-11 (SP6-11). These simulations were initiated from the cryo-EM structure of SP-bound NK1R, with the intracellular G protein removed and, for SP6-11-bound simulations, residues 1-5 of SP removed.

The peptide-bound NK1R structure was prepared for simulation with Maestro (Schrödinger Release 2019-4: Maestro, Schrödinger, LLC, New York, NY, 2019). Missing amino acid side chains were modeled using Prime^74,75^. Residues 226-237 are missing in the cryo-EM structure and were not modeled in. Neutral acetyl and methylamide groups were added to cap the N- and C-termini, respectively, of the NK1R protein chains. The N-termini of SP and SP6-11 were prepared in their charged form, while the C-termini were amidated. Titratable residues were kept in their dominant protonation state at pH 7, except for E2.50 (E78) and D3.49 (D129), which were protonated to their neutral form, as studies indicate that these conserved residues are protonated in active-state class-A GPCRs^76,77^. Histidine residues were modeled as neutral, with a hydrogen atom bound to either the delta or epsilon nitrogen depending on which tautomeric state optimized the local hydrogen-bonding network. Dowser^78^ was used to add water molecules to protein cavities, and the protein structures were aligned on the transmembrane (TM) helices of the inactive NK1R crystal structure (PDB ID: 6HLP)^38^ in the Orientation of Proteins in Membranes (OPM) database^79^. The aligned structures were inserted into a pre-equilibrated palmitoyloleoyl-phosphatidylcholine (POPC) membrane bilayer using Dabble^80^. Sodium and chloride ions were added to neutralize each system at a concentration of 150 mM. The final systems comprised 57605-58494 atoms, including 134 lipid molecules and 11565-11831 water molecules. Approximate system dimensions were 80 Å x 80 Å x 94 Å.

### Molecular Dynamics: Simulation protocols

For each simulation condition (SP-bound and SP6-11-bound), we performed 12 independent simulations (~2-μs each) in which initial atom velocities were assigned randomly and independently. We employed the CHARMM36m force field for protein molecules, the CHARMM36 parameter set for lipid molecules and salt ions, and the associated CHARMM TIP3P model for water^81–83^. Simulations were run using the AMBER18 software^84^ under periodic boundary conditions with the Compute Unified Device Architecture (CUDA) version of Particle-Mesh Ewald Molecular Dynamics (PMEMD) on one GPU^85^.

After energy minimization, the systems were first heated over 12.5 ps from 0 K to 100 K in the NVT ensemble using a Langevin thermostat with harmonic restraints of 10.0 kcal·mol^-1^·Â^-2^ on the non-hydrogen atoms of the lipids, protein, and ligand. Initial velocities were sampled from a Boltzmann distribution. The systems were then heated to 310 K over 125 ps in the NPT ensemble. Equilibration was performed at 310 K and 1 bar in the NPT ensemble, with harmonic restraints on the protein and ligand non-hydrogen atoms tapered off by 1.0 kcal·mol^-1^·Â^-2^ starting at 5.0 kcal·mol^-1^·Â^-2^ in a stepwise manner every 2 ns for 10 ns, and finally by 0.1 kcal·mol^-1^·Â^-2^ every 2 ns for an additional 18 ns. All restraints were completely removed during production simulation. Production simulations were performed at 310 K and 1 bar in the NPT ensemble using the Langevin thermostat and Monte Carlo barostat. The simulations were performed using a timestep of 4.0 fs while employing hydrogen mass repartitioning^86^. Lengths of bonds to hydrogen atoms were constrained using SHAKE^87^. Non-bonded interactions were cut off at 9.0 Å, and long-range electrostatic interactions were calculated using the particle-mesh Ewald (PME) method with an Ewald coefficient (β) of approximately 0.31 Å and B-spline interpolation of order 4. The PME grid size was chosen such that the width of a grid cell was approximately 1 Å.

### Molecular Dynamics: Simulation analysis protocols

The AmberTools17 CPPTRAJ package^88^ was used to reimage trajectories, while Visual Molecular Dynamics (VMD)^89^ and PyMOL (The PyMOL Molecular Graphics System, Version 2.0 Schrodinger, LLC.) were used for visualization. MDAnalysis^90^ was used for simulation analysis.

Root mean square fluctuation (RMSF) values shown in Figure 4B measure the extent to which a group of atoms fluctuates around its average position in simulation and is thus a measure for mobility. The first 500 ns of each simulation trajectory were omitted from this analysis to avoid including any initial relaxation of the system in the measurement. The analysis was performed on 1501 frames per simulation, where each frame is separated by 1 ns. For each simulation, an average position of each atom in a specified group (residues 6-11, F7, or M11; for each of these three groups all atoms were included) was calculated. Then, the RMSF was obtained as the time-average of the RMSD to the average structure for each simulation. For the RMSF of residues 6-11, trajectories were aligned to the initial cryo-EM structure on all transmembrane helix Cα atoms. For the RMSF of residues F7 and M11, trajectories were aligned to the initial cryo-EM structure on all Cα atoms of residues 6-11 of SP to better capture the individual residue movement independent of the overall movement of the entire peptide. Each bar in Figure 4 represents the mean RMSF value over all 12 simulations per condition (either SP or SP6-11) and the error bars denote the standard error of the mean. To test statistical significance, we performed two-sided t-tests of unequal variance (Welch’s t-tests).

For each of the 3 renderings in Figure 4A and for each of the 3 renderings in Figure 4C, we chose 10 representative simulation frames illustrating the dynamics of the backbone of residues 6-11 and the side chains of F7 and M11.

## Data Availability

All data generated or analyzed during this study are included in this published article and its Supplementary Information. Coordinates for SP-NK1R-miniG_s/q70_, SP-NK1R-miniG_s399_, and the SP6-11-NK1R-miniG_s/q70_ complex have been deposited in the Protein Data Bank under accession codes XXXX, XXXX, XXXX, respectively. Unsharpened cryo-EM maps for SP-NK1R-miniG_s/q70_, SP-NK1R-miniG_s399_, and the SP6-11-NK1R-miniG_s/q70_ complex have been deposited in the Electron Microscopy Data Bank under accession codes XXXXX, XXXXX, and XXXXX, respectively. Sharpened cryo-EM maps for SP-NK1R-miniG_s/q70_, SP-NK1R-miniG_s399_, and the SP6-11-NK1R-miniG_s/q70_ complex have been deposited in the Electron Microscopy Data Bank under accession codes XXXXX, XXXXX, and XXXXX, respectively.

## Acknowledgements

This work was supported by National Institutes of Health (NIH) grants DP5OD023048 (A.M.), R01GM127359 (R.O.D.), and a National Health and Medical Research Council (NHMRC) Project Grant APP1138448 (D.M.T), NHMRC Investigator Grant APP1196951 (D.M.T), Australian Research Council DECRA grant DE170100152 (D.M.T.), and Australian Research Council Centre of Excellence in Convergent Bio-Nano Science and Technology (N.A.V). This material is based upon work supported by the National Science Foundation Graduate Research Fellowship Program (J.A.H.) under Grant No. 2034836. Any opinions, findings, and conclusions or recommendations expressed in this material are those of the author(s) and do not necessarily reflect the views of the National Science Foundation. C.-M.S. was funded by the Human Frontier Science Program (LT000916/2018-L). Cryo-EM equipment at UCSF is partially supported by NIH grants S10OD020054 and S10OD021741. Some of this work was performed at the Stanford-SLAC Cryo-EM Center (S2C2), which is supported by the National Institutes of Health Common Fund Transformative High-Resolution Cryo-Electron Microscopy program (U24 GM129541). The content is solely the responsibility of the authors and does not necessarily represent the official views of the National Institutes of Health. Y.C. is an Investigator of Howard Hughes Medical Institute. A.M. acknowledges support from the Pew Charitable Trusts, the Esther and A. & Joseph Klingenstein Fund and the Searle Scholars Program.

## Author Contributions

J.A.H. purified NK1R-constructs, Gβ_1_γ_2_, and Nb35, established biochemical approaches to reconstitute a NK1R-miniG protein complex, and generated all NK1R mutants. B.F. purified Gβ_1_γ_2_, and Nb35, prepared samples for cryo-EM, identified optimal freezing conditions for cryo-EM, screened samples by cryo-EM, collected cryo-EM data, and determined high resolution cryo-EM maps by extensive image processing under the guidance of A.M. and Y.C.. J.A.H. built and refined models of NK1R-miniG protein complexes with input from B.F. and A.M.. A.B.G. generated NK1R stable cell lines and performed cellular signaling experiments under the guidance of N.A.V. and D.M.T.. M.A.D. and C.-M.S. performed and analyzed molecular dynamics simulations under the guidance of R.O.D. The manuscript was written by J.A.H., B.F., A.B.G., M.A.D., C.-M.S., R.O.D, and A.M., with edits from N.A.V., D.M.T., and Y.C. and with approval from all authors. The overall project was supervised by A.M.

