## Supplementary_Information for "Selective G protein signaling driven by Substance P-Neurokinin Receptor structural dynamics"

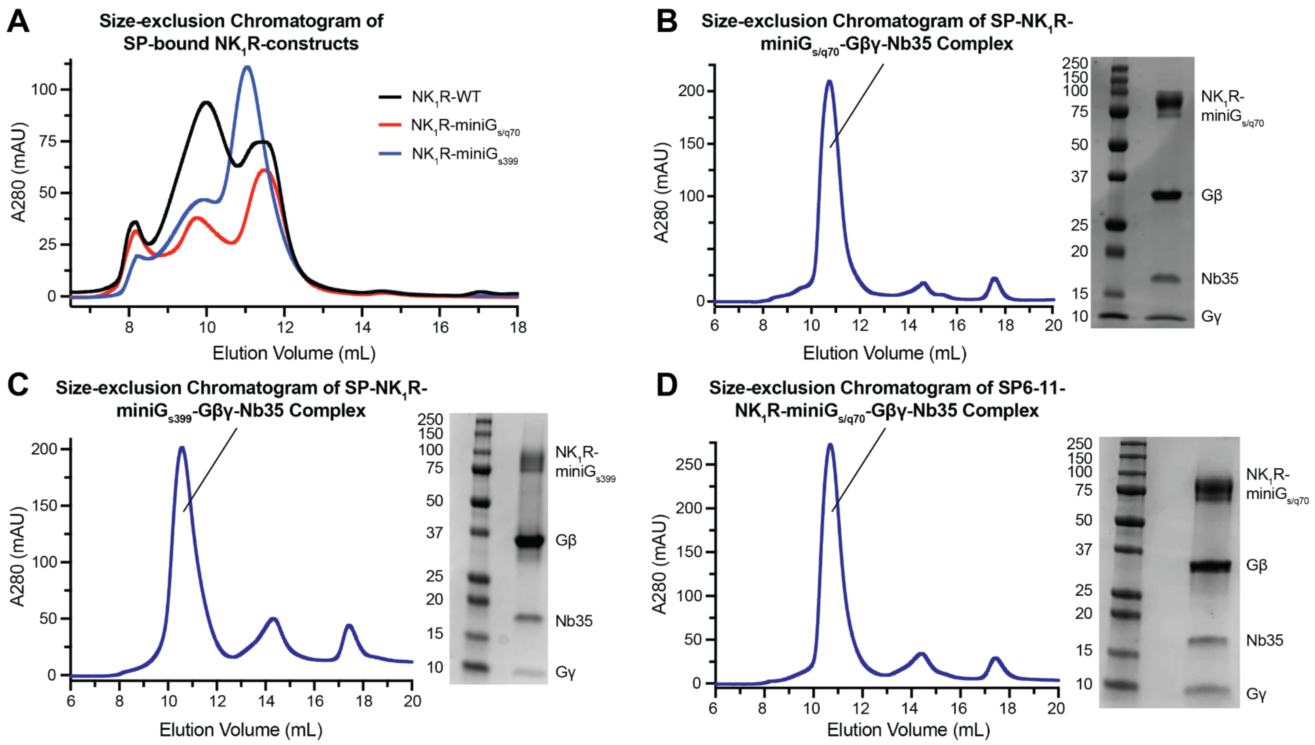


**Supplementary Figure 1. Biochemistry of active-state NK1R-miniG protein complexes. (A)** Size-exclusion chromatography of SP-bound NK1R and NK1R-miniG$_{s/q70}$ fusion shows an increase in the monomeric species for NK1R-miniG fusion proteins. Size-exclusion chromatography traces and SDS-PAGE gels of purified SP-bound NK1R-miniG$_{s/q70}$ complex (**B**), SP-bound NK1R-miniG$_{s399}$ complex (**C**), and SP6-11-bound NK1R-miniG$_{s/q70}$ complex (**D**).

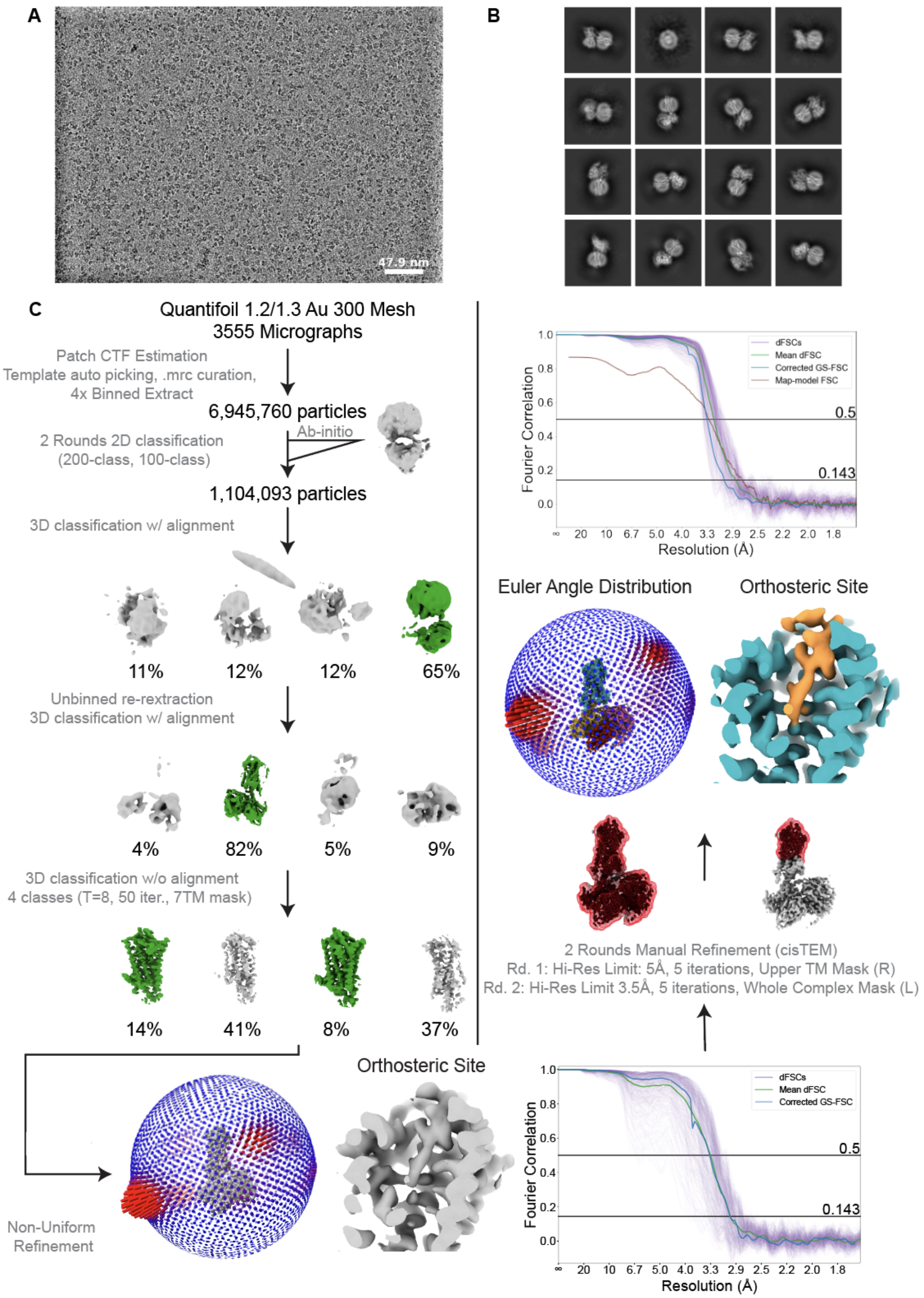


**Supplementary Figure 2. CryoEM data processing workflow for SP-NK1R-miniG$_{s/q70}$ heterotrimeric complex.**

Representative micrograph (**A**) and 2D-class averages (**B**) for SP-NK1R-miniG$_{s/q70}$ complex. (**C**) A flowchart representation of the processing pipeline used for structural determination of the

SP-NK1R-miniG$_{s/q70}$ complex. Contrast transfer function (CTF) estimation, 2D classification and all 3D classification jobs with alignment were performed with cryoSPARC. 3D classification without alignment was performed with RELION using a mask encompassing only the receptor transmembrane and final focused refinements were performed with cisTEM. Focused refinement masks are shown as red mesh. Gold-standard fourier shell correlation (GS-FSC) was calculated from a cryoSPARC Local Resolution job using the focused refinement mask encompassing the entire SP-NK1R-miniG$_{s/q70}$ complex. A viewing distribution plot was generated using scripts from the pyEM software suite and visualized in ChimeraX. Directional FSC curves (dFSCs) are shown as purple lines and were determined as previously described in Dang, S. et al[67].

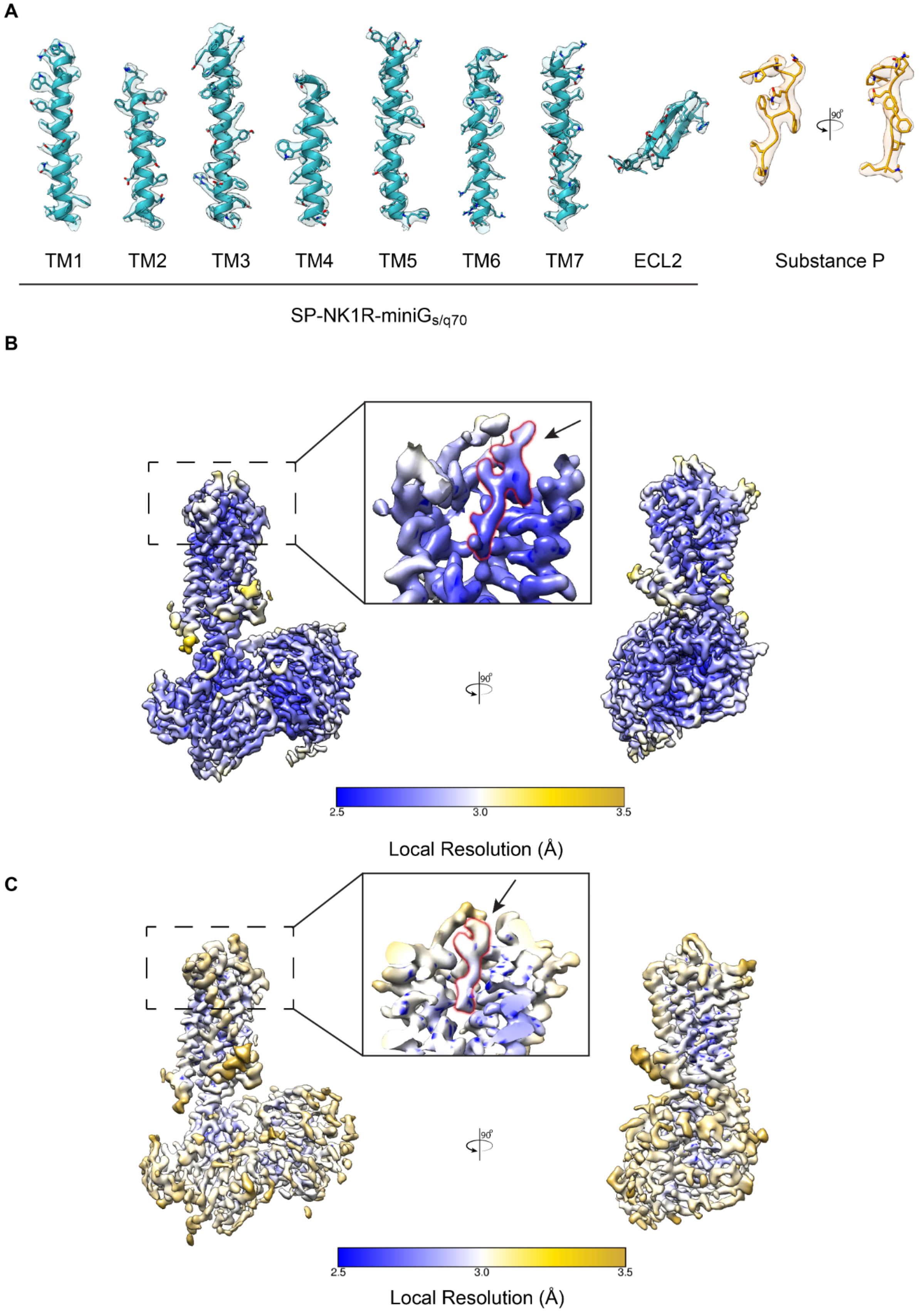
A
TM1
TM2
TM3
TM4
TM5
TM6
TM7
ECL2
Substance P
90°
SP-NK1R-miniGs/q70
B
90°
2.5
3.0
3.5
Local Resolution (Å)
C
90°
2.5
3.0
3.5
Local Resolution (Å)

**Supplementary Figure 3. Cryo-EM density map for NK1R-miniG$_{s/q70}$ heterotrimeric complex.**

(**A**) Unsharpened Cryo-EM density map for individual NK1R helices and Substance P density as determined by extending a 2.5 Å radius away from each modeled atom. Local resolution estimation of unsharpened Cryo-EM density maps for (**B**) SP-NK1R-miniG$_{s/q70}$ and (**C**) SP6-11-NK1R-miniG$_{s/q70}$ heterotrimeric complex from cryoSPARC. SP and SP6-11 density are highlighted in and shown at equivalent enclosed volume thresholds.

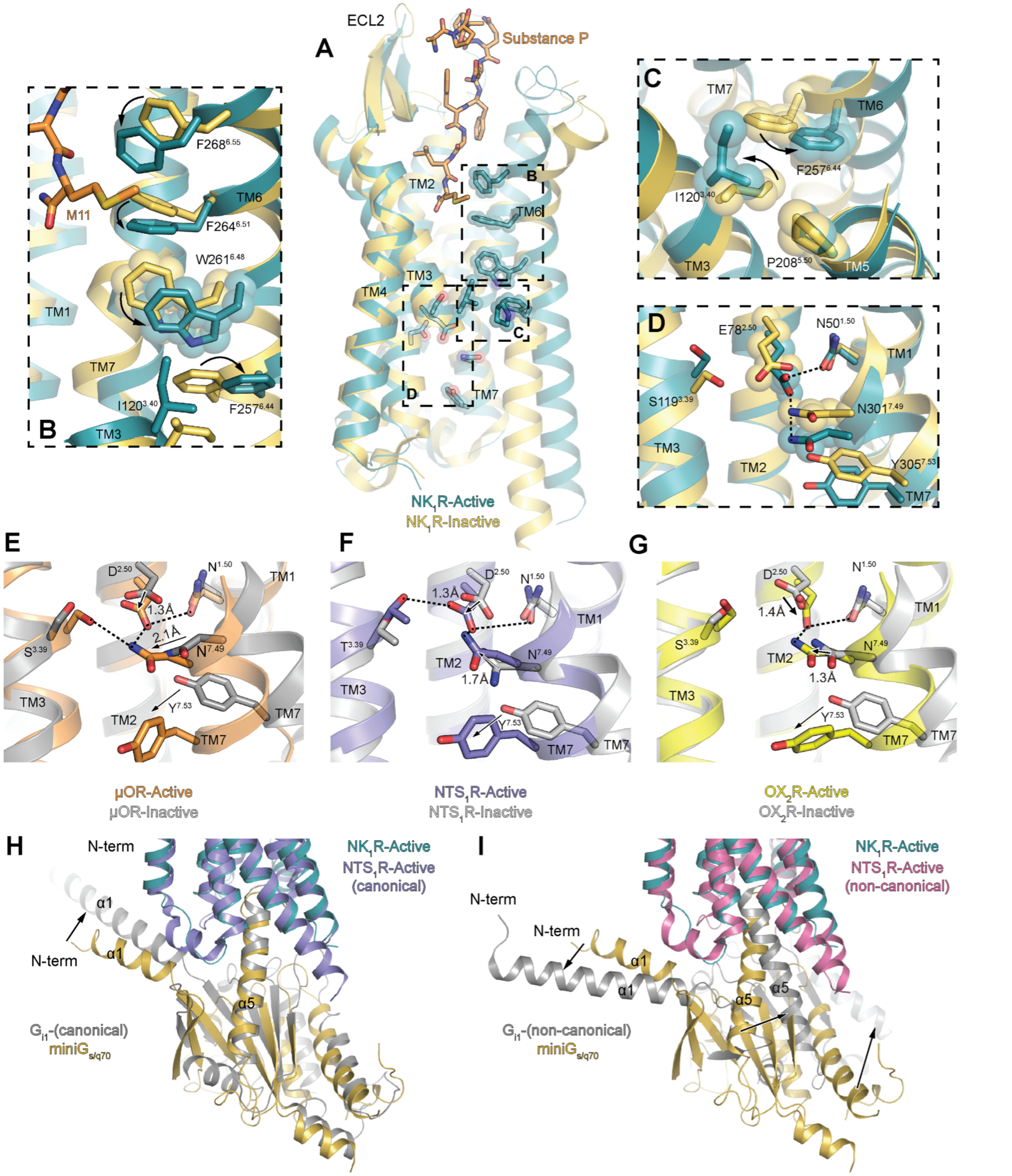


**Supplementary Figure 4. Structural hallmarks of NK1R activation.**

(**A**) Alignment of the SP-NK1R-miniG$_{s/q70}$ structure with an inactive-state NK1R structure (PDB: 6HLP[38]) reveals rearrangement of NK1R structural motifs indicative of class A GPCR activation, including: (**B**) displacement of the $W^{6.48}$ 'toggle-switch' and (**C**) rearrangement of the '$P^{5.50}I^{3.40}F^{6.44}$' connector motif. (**D**) The non-canonical $E78^{2.50}$-$N301^{7.49}$ interaction in NK1R is unchanged between inactive- and active-state structures. We compared the NK1R $E78^{2.50}$-$N301^{7.49}$

interaction to the $D^{2.50}$-$N^{7.49}$ interaction in three class A neuropeptide-binding GPCRs, including: (**E**) the μ-opioid receptor (Active PDB: 5C1M[29], Inactive PDB: 4DKL[58]), (**F**) the neurotensin 1 receptor (Active PDB: 6OS9[35], Inactive PDB: 4BUO[59]), and (**G**) the orexin 2 receptor (Active PDB: 7L1U, Inactive PDB: 5WQC). Alignment of the SP-NK1R-mini$G_{s/q70}$ structure with (**H)** canonical (PDB: 6OS9[35]) and (**I**) 'non-canonical' (PDB: 6OSA[35]) active-state $NTS_1R$ reveals that the mini$G_{s/q70}$ protein adopts the canonical G protein coupling orientation.

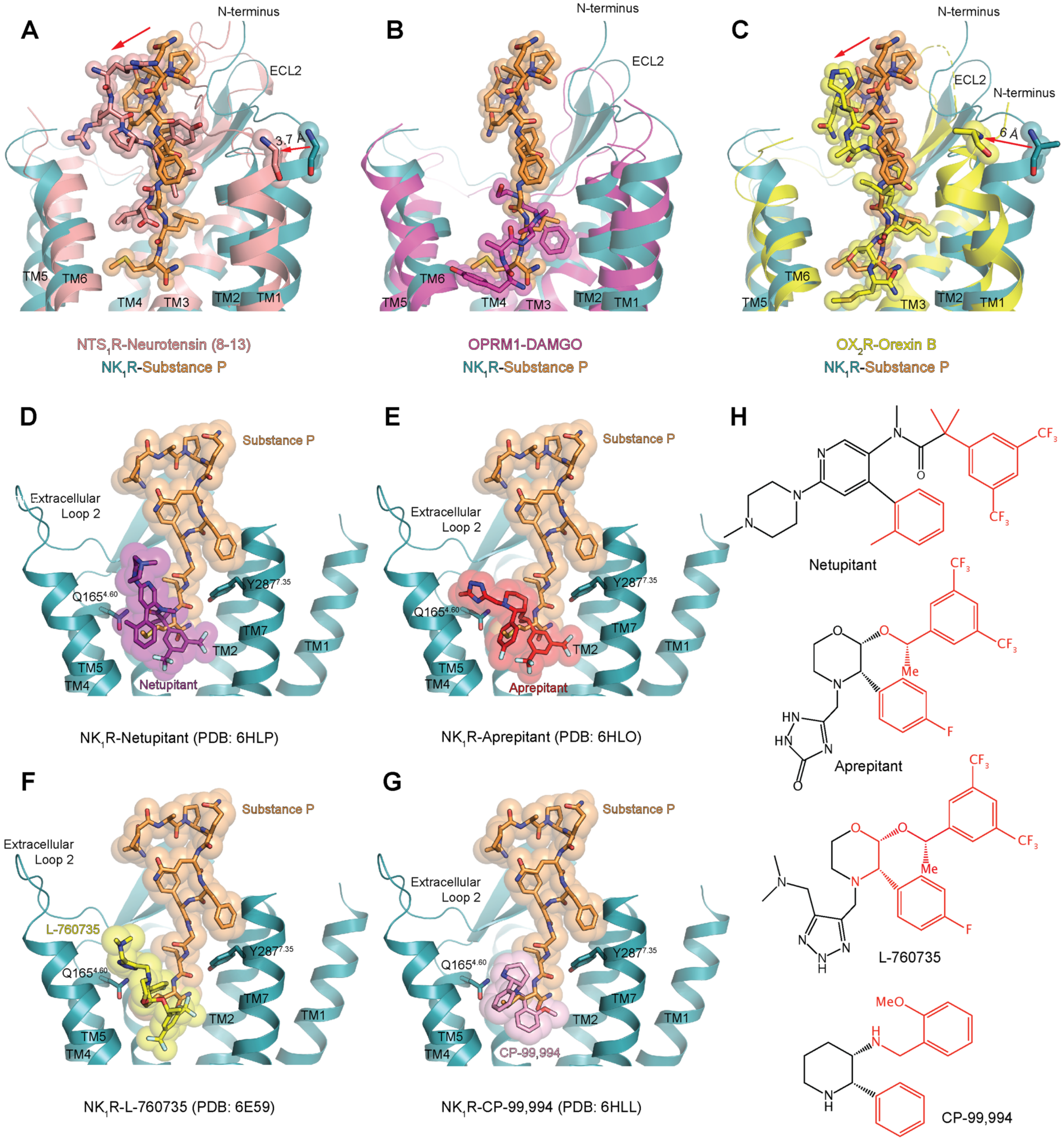


**Supplementary Figure 5. Comparison of SP-NK1R binding site to related Neuropeptide GPCRs and Inactive-State NK1R Structures.**

Comparison of SP-bound NK1R-miniG$_{s/q70}$ structure to neuropeptide GPCRs bound to peptidergic ligands, including: **A**) the neurotensin 1 receptor bound to neurotensin 8-13 (PDB: 4GRV[40]), **B**) the μ-opioid receptor bound to the peptide mimetic agonist DAMGO (PDB: 6DDE[41]), and **C**) the orexin 2 receptor bound to orexin B (PDB: 7L1U[42]). Alignment of SP-bound NK1R with inactive-state NK1R structures, including: (**D**) netupitant-bound (PDB: 6HLP[38]), (**E**) aprepitant-bound (PDB: 6HLO[38]), (**F**) L-760,735-bound (PDB: 6E59[43]), and (**G**) CP-99,994-bound NK1R (PDB: 6HLL[38]). (**H**) Antagonist chemical structures shown with regions that compete with SP binding site in red.

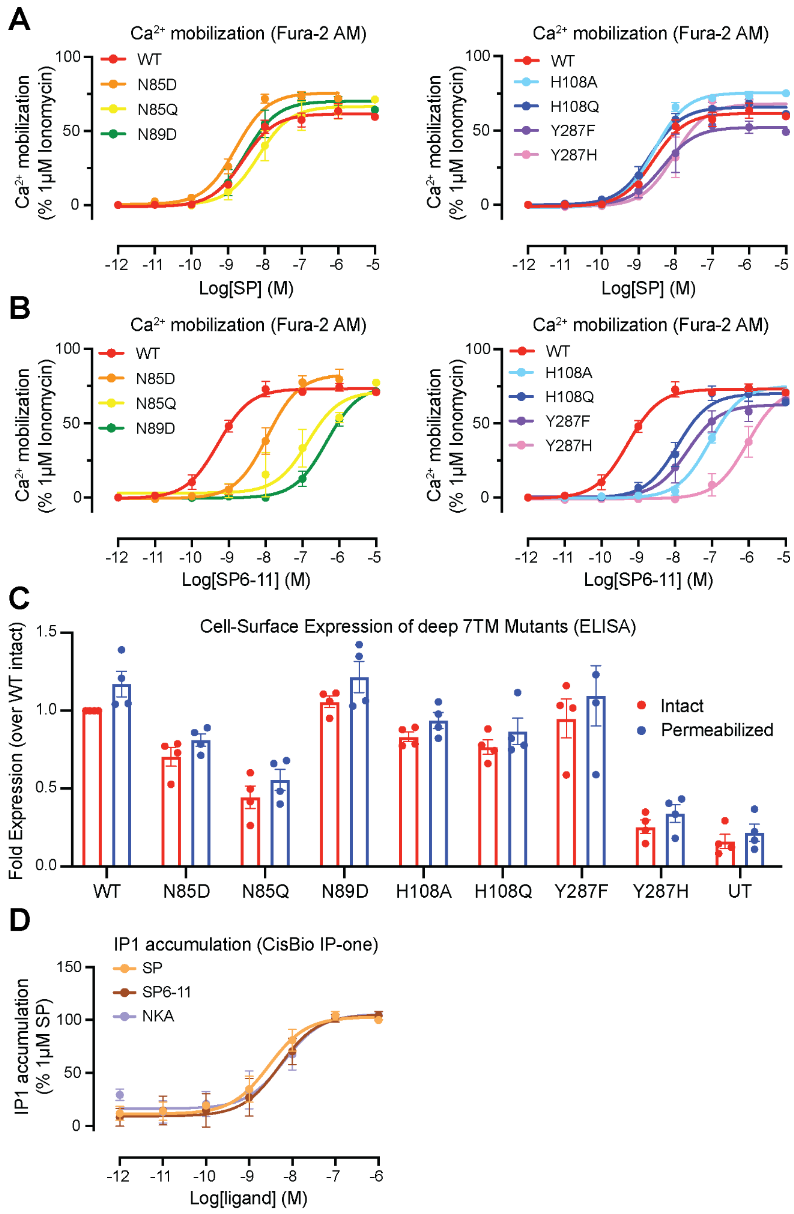


**Supplementary Figure 6. Signaling studies for NK1R mutations in the deep 7TM region.** $Ca^{2+}$ mobilization of wild-type and NK1R mutants after stimulation with (**A**) SP and (**B**) SP6-11. Signaling graphs represent the global fit of grouped data ± s.e.m. from n $\geq$ 3 independent biological replicates. Full quantitative parameters from this experiment are listed in Supplementary Table 3. (**C**) Cell-surface expression of deep 7TM NK1R mutants as determined by ELISA. Untransfected (UT) control shows low ELISA signal. Bar graphs represent mean ± s.e.m. from n = 4 independent biological replicates. (**D**) IP1 accumulation of wild-type NK1R after stimulation with SP, NKA, and N-terminally truncated SP analogs. Signaling graphs represent the global fit ± s.e.m. from n = 3 independent biological replicates. Full quantitative parameters from this experiment are listed in Supplementary Table 2.

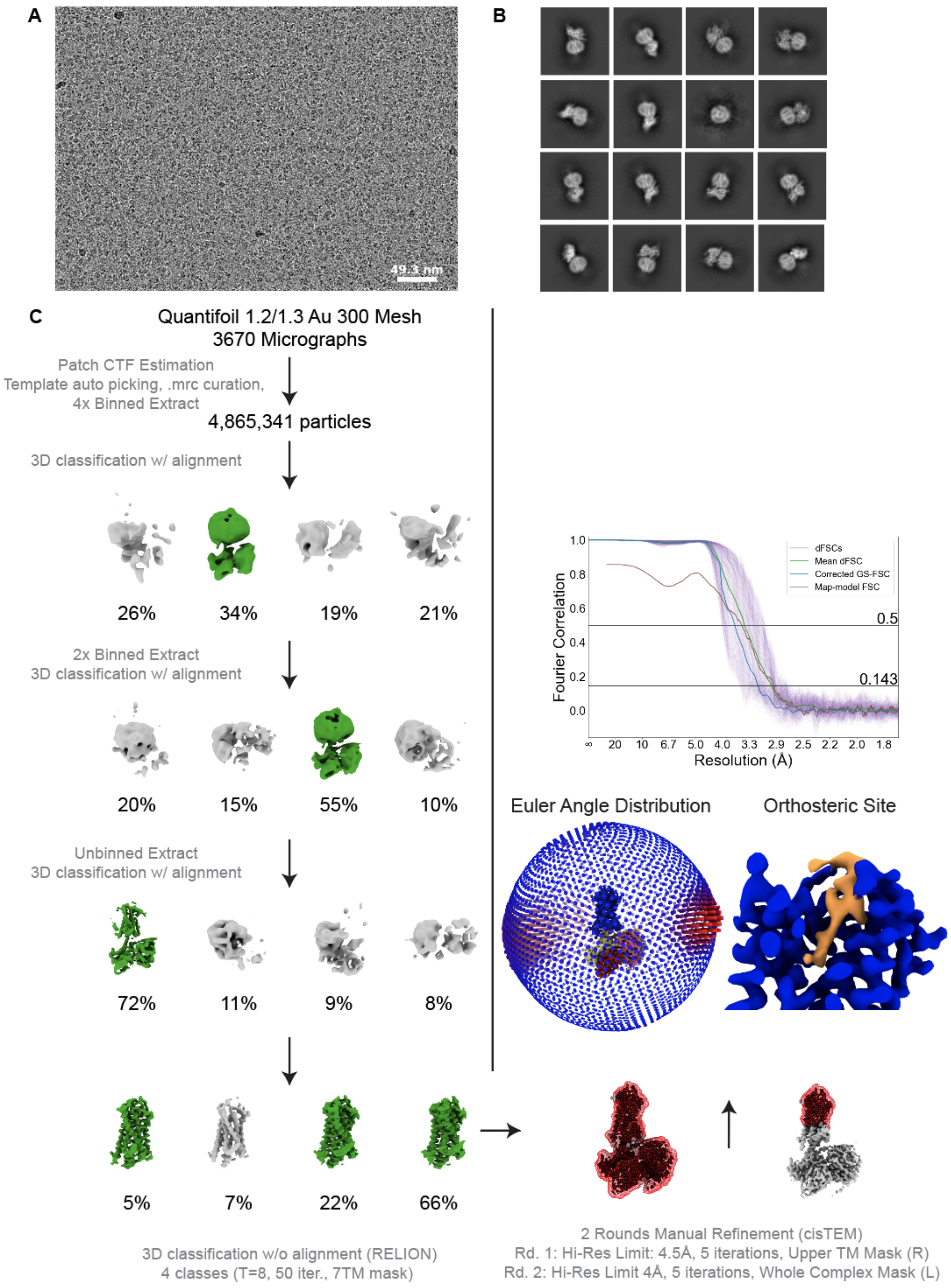


**Supplementary Figure 7**. **CryoEM data processing for SP-NK1R-miniG$_{s399}$ complex.** Representative micrograph (**A**) and 2D-class averages (**B**) for SP-NK1R-miniG$_{s399}$ complex. (**C**) A flowchart representation of the processing pipeline used for structural determination of the SP-NK1R-miniG$_{s399}$ complex. CTF Estimation, 2D classification and all 3D classification jobs with alignment were performed with cryoSPARC. 3D classification without alignment was performed with RELION using a mask encompassing only the receptor transmembrane and final focused refinements were performed with cisTEM. Focused refinement masks are shown as red mesh. Gold-standard fourier shell correlation (GS-FSC) was calculated from a cryoSPARC Local Resolution job using the focused refinement mask encompassing the entire SP-NK1R-miniG$_{s399}$ complex. A viewing distribution plot was generated using scripts from the pyEM software suite and visualized in ChimeraX. Directional FSC (dFSC) are shown as purple lines and were determined as previously described[67].

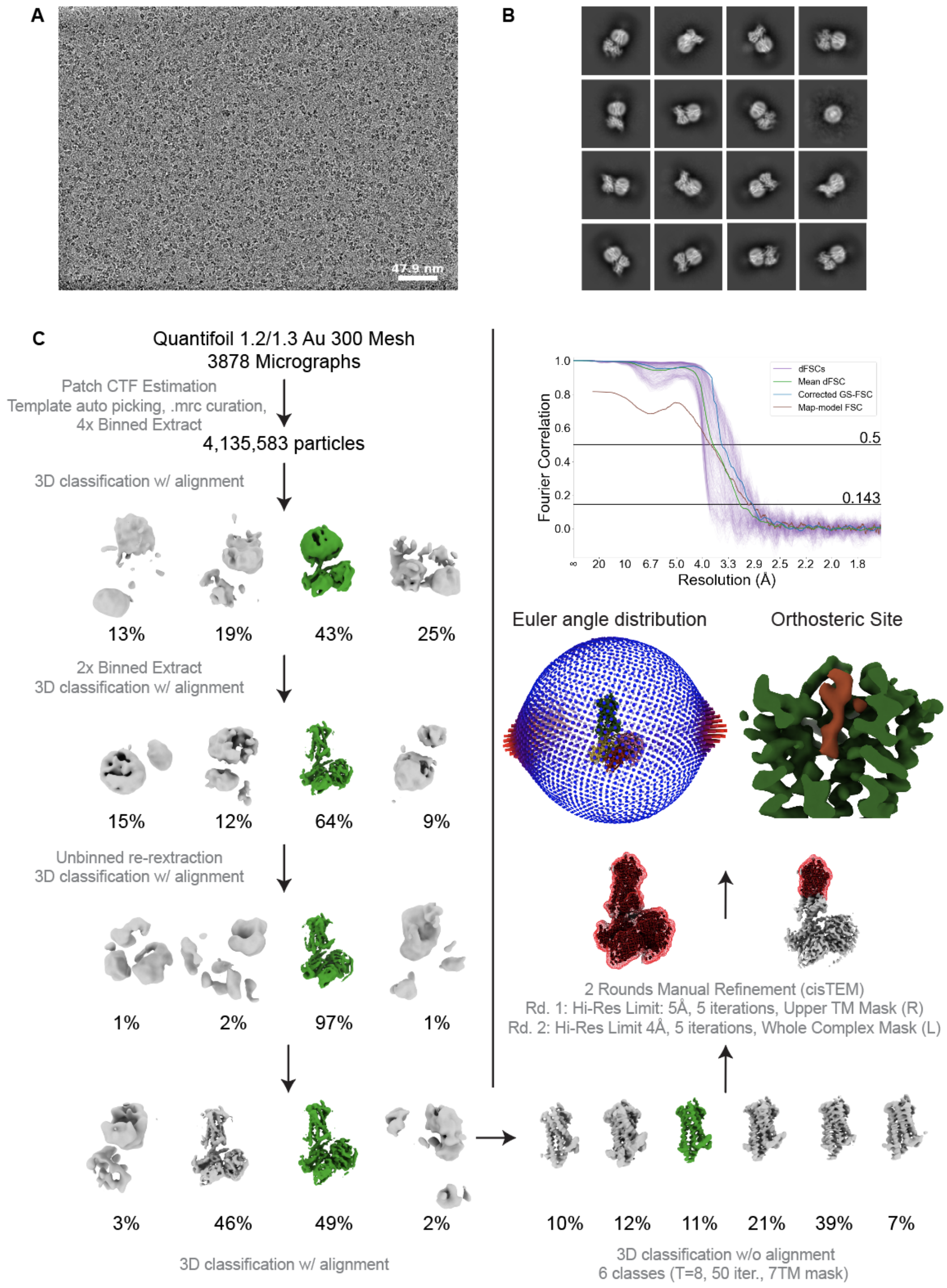


**Supplementary Figure 8. CryoEM data processing for SP6-11-NK1R-miniG$_{s/q70}$ complex.** Representative micrograph (**A**) and 2D-class averages (**B**) for SP6-11-NK1R-miniG$_{s/q70}$ complex. (**C**) A flowchart representation of the processing pipeline used for structural determination of the SP6-11-NK1R-miniG$_{s/q70}$ complex. CTF Estimation, 2D classification and all 3D classification jobs with alignment were performed with cryoSPARC. 3D classification without alignment was performed with RELION using a mask encompassing only the receptor transmembrane and final focused refinements were performed with cisTEM. Focused refinement masks are shown as red mesh. GS-FSC was calculated from a cryoSPARC Local Resolution job using the focused refinement mask encompassing the entire SP6-11-NK1R-miniG$_{s/q70}$ complex. A viewing distribution plot was generated using scripts from the pyEM software suite and visualized in ChimeraX. Directional FSC curves (dFSC) are shown in purple lines and were determined as previously described[67].

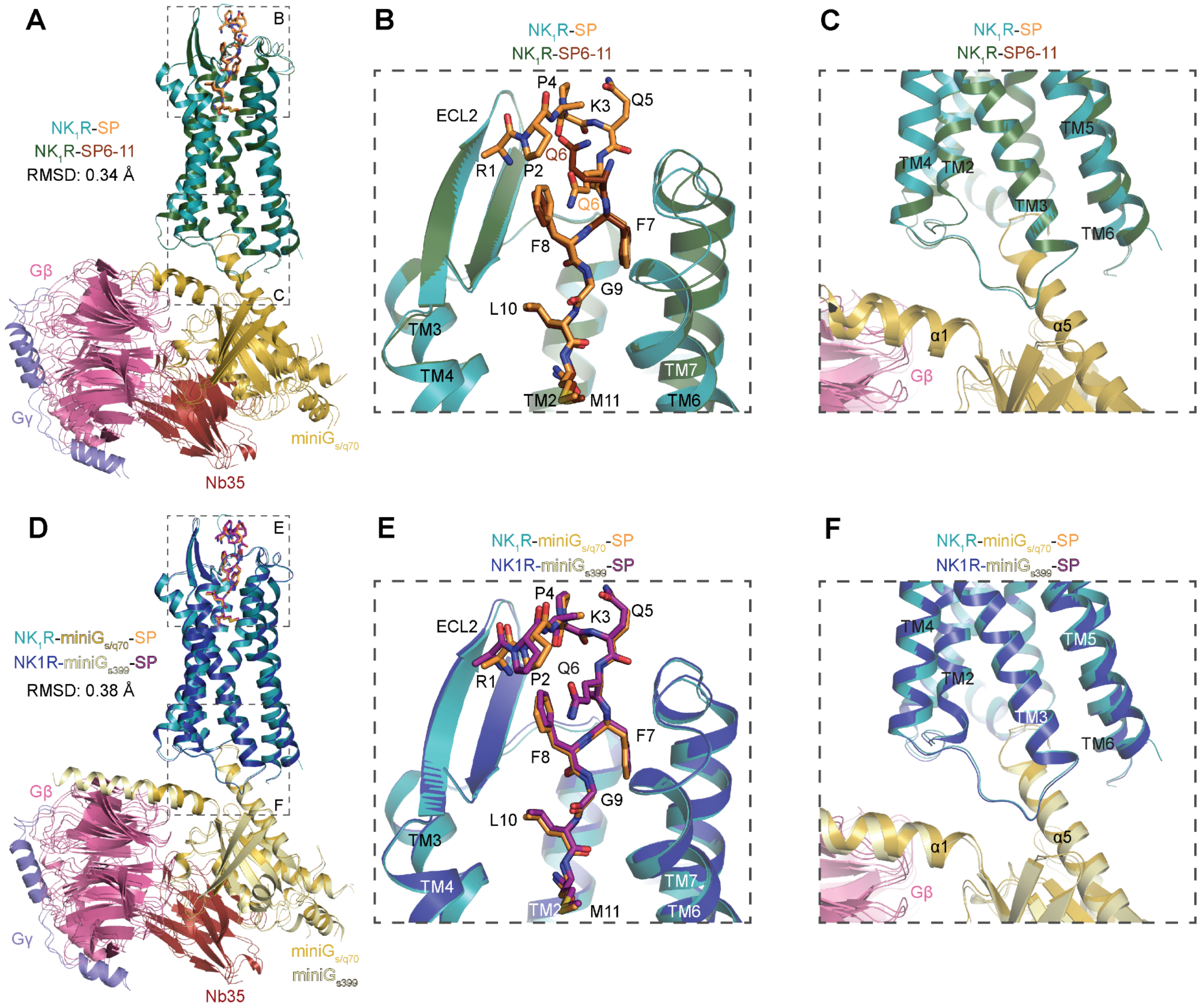


**Supplementary Figure 9. Comparison of SP- and SP6-11-bound NK1R G protein-complexes**

Alignment of SP-NK1R-miniG$_{s/q70}$ and SP6-11-NK1R-miniG$_{s/q70}$ through NK1R 7TM domain reveals minimal changes in (**A**) overall 7TM architecture, (**B**) overall peptide binding poses, and (**C**) insertion of miniG protein α5 helix in NK1R core. Alignment of SP-NK1R-miniG$_{s/q70}$ and SP-NK1R-miniG$_{s399}$ through NK1R 7TM domain reveals minimal changes in (**D**) overall 7TM architecture, (**E**) overall Substance P binding pose, and (**F**) insertion of miniG protein α5 helix in NK1R core.

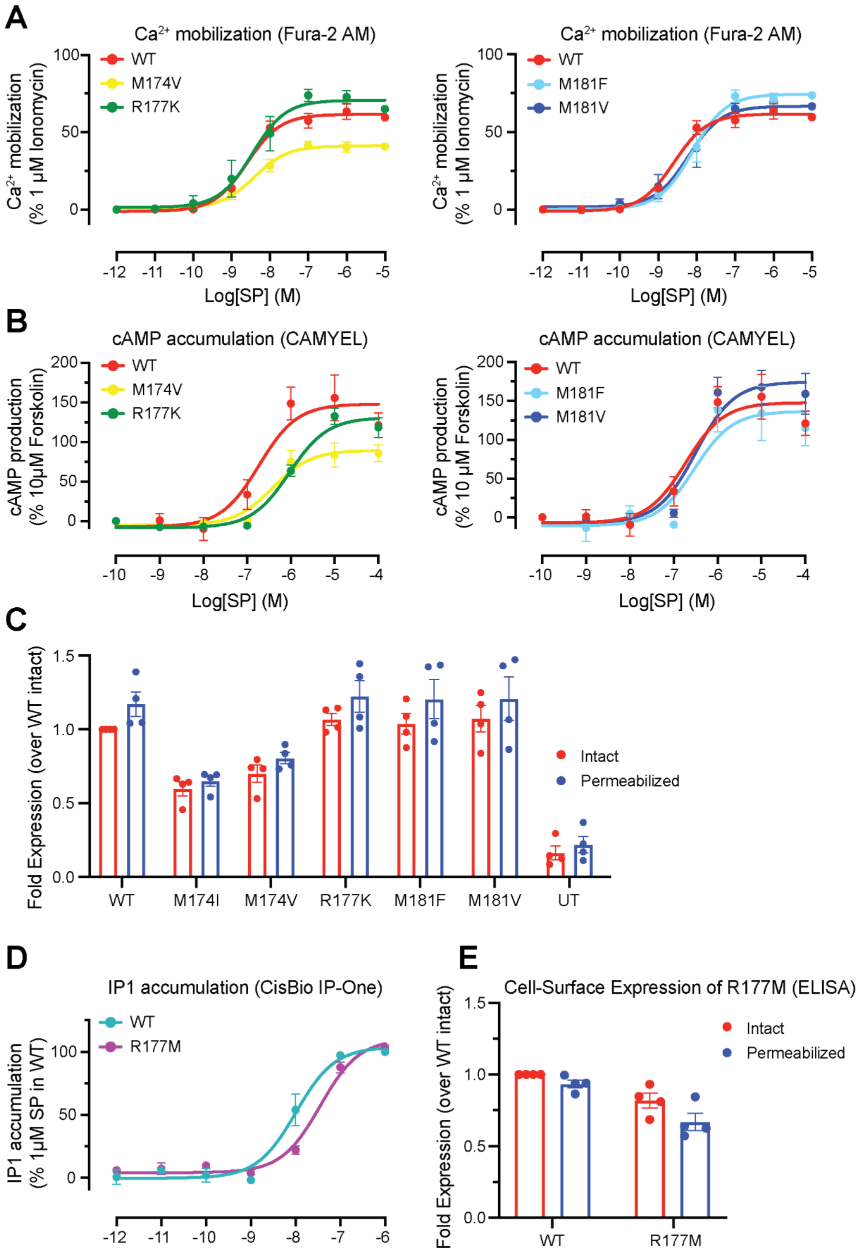


**Supplementary Figure 10. Signaling studies for NK1R ECL2 mutations.**
(**A,B**) $Ca^{2+}$ mobilization and cAMP accumulation of wild-type and ECL2 NK1R mutants after stimulation with SP. Signaling graphs represent the global fit of grouped data ± s.e.m. from n $\geq$ 3 independent biological replicates. Full quantitative parameters from this experiment are listed in Supplementary Table 3. (**C**) Cell-surface expression of ECL2 NK1R mutants as determined by ELISA. Untransfected (UT) control shows low ELISA signal. Bar graphs represent mean ± s.e.m. from n = 4 independent biological replicates. (**D**) IP1 accumulation of wild-type and R177M NK1R after stimulation with SP. Signaling graphs represent the global fit of grouped data ± s.e.m. from n = 4 independent biological replicates. Full quantitative parameters from this experiment are

listed in Supplementary Table 3. (**E**) Cell-surface expression of R177M NK1R mutant as determined by ELISA. Bar graphs represent mean ± s.e.m. from n = 4 independent biological replicates.

**Supplementary Table 1. Cryo-EM data collection and model refinement statistics.**

| **Sample:** | SP-NK1R-miniG$_{s/q70}$ Gβγ-Nb35 | SP6-11-NK1R-miniG$_{s/q70}$ Gβγ-Nb35 | SP-NK1R-miniG$_{s399}$ Gβγ-Nb35 |
|---|---|---|---|
| **Unsharpened map EMDB:** | XXXXX | XXXXX | XXXXX |
| **Sharpened map EMDB:** | XXXXX | XXXXX | XXXXX |
| **PDB:** | XXXX | XXXX | XXXX |
| **Data Collection & Processing** | | | |
| Microscope/Detector | Titan Krios/Gatan K3 with Gatan Bioquantum Energy Filter | | |
| Imaging Software and collection | Serial EM, 3x3 image Shift | | EPU 2.1, AFIS |
| Magnification | 105,000 | | 105,000 |
| Voltage (kV) | 300 | | 300 |
| Electron exposure (e-/Å$^2$) | 67 | | 49 |
| Dose rate (e-/pix/sec) | 8 | | 24 |
| Frame exposure (e-/Å$^2$) | 0.55 | | 0.82 |
| Defocus range (µm) | -0.8 to -2.0 | | -0.8 to -2.1 |
| Pixel size (Å) | 0.835 (physical) | | 0.860 (physical) |
| Micrographs | 3755 | 3878 | 3670 |
| **Reconstruction** | | | |
| Autopicked particles (template-based in cryosparc) | 6,945,760 | 4,135,583 | 4,865,341 |
| Particles in final refinement | 122,220 | 59,926 | 288,659 |
| Symmetry imposed | C1 | C1 | C1 |
| Map sharpening *B* factor (Å$^2$) | unsharpened or -87 | unsharpened or -80 | unsharpened or -120 |
| Map resolution, global FSC (Å) | | | |
| FSC 0.5, unmasked/masked | 3.5/3.3 | 4.0/3.8 | 3.9/3.5 |
| FSC 0.143, unmasked/masked | 3.2/3.0 | 3.6/3.2 | 3.3/3.1 |
| **Refinement** | | | |
| Initial model used (PDB codes) | 6HLP, 6LI3, 3SN6 | SP-NK1R-miniG$_{s/q70}$ | SP-NK1R-miniG$_{s/q70}$ |
| Model resolution (Å) | | | |
| FSC 0.5 unmasked/masked | 3.4/3.2 | 3.9/3.7 | 3.6/3.3 |
| Model composition | | | |
| Non-hydrogen atoms | 7316 | 7202 | 7434 |
| Protein residues | 968 | 960 | 986 |
| Ligands | N: 1 | N: 1 | N: 1 |
| *B* Factors (Å$^2$) | | | |
| Protein | 35.99 | 50.1 | 74.3 |
| Ligand | 30.0 | 29.3 | 57.3 |
| R.m.s. deviations | | | |
| Bond lengths (Å) | 0.004 | 0.004 | 0.004 |
| Bond angles (°) | 0.815 | 0.842 | 0.746 |
| Validation | | | |
| Molprobity score | 1.27 | 1.43 | 1.31 |
| Clashscore | 3.58 | 4.29 | 2.49 |
| Poor rotamers (%) | 0 | 0 | 0 |
| EM ringer score | 4.52 | 2.73 | 3.42 |
| CaBLAM outliers (%) | 1.31 | 1.87 | 1.72 |
| Ramachandran Plot | | | |
| Favored (%) | 97.35 | 96.57 | 95.93 |
| Allowed (%) | 2.65 | 3.43 | 4.07 |
| Outliers (%) | 0 | 0 | 0 |

**Supplementary Table 2. Summary of tachykinin signaling studies.** Values are expressed as mean $pEC_{50}$ or $E_{max}$ ± s.e.m. from (n) independently fit biological replicates. Mean $pEC_{50}$ and $E_{max}$ values for NKA and SP6-11 are compared in a one-way analysis of variance (ANOVA) with Dunnett's multiple comparison–corrected post hoc test against SP represented by * = p $\leq$ 0.033, ** = p $\leq$ 0.002.

| | Ligand | $Ca^{2+}$ $pEC_{50}$ | $Ca^{2+}$ $E_{max}$ | cAMP $pEC_{50}$ | cAMP $E_{max}$ | IPone $pEC_{50}$ | IPone Emax |
|---|---|---|---|---|---|---|---|
| NK1R | SP | 8.7 ± 0.2 (10) | 97 ± 8 (10) | 6.9 ± 0.2 (12) | 122 ± 11 (12) | 8.5 ± 0.3 (3) | 90 ± 5 (3) |
| | NKA | 8.7 ± 0.4 (6) | 89 ± 10 (6) | 6.1 ± 0.1 (7) * | 111 ± 11 (7) | 8.3 ± 0.5 (3) | 86 ± 6 (3) |
| | SP6-11 | 8.9 ± 0.5 (4) | 96 ± 8 (4) | 5.7 ± 0.1 (4) ** | 129 ± 17 (4) | 8.4 ± 0.3 (3) | 92 ± 9 (3) |

**Supplementary Table 3. Summary of mutant NK1R signaling studies.** Values are expressed as mean $pEC_{50}$ or mean $E_{max}$ ± s.e.m. from (n) independently fit biological replicates. Mean $pEC_{50}$ and $E_{max}$ values for NK1R mutants stimulated with SP are compared in a one-way analysis of variance (ANOVA) with Dunnett's multiple comparison–corrected post hoc test against wild-type NK1R stimulated with SP, (* = p ≤ 0.033, ** = p ≤ 0.002, *** = p ≤ 0.0002, **** = p ≤ 0.0001). Mean $pEC_{50}$ and $E_{max}$ values for NK1R mutants stimulated with SP6-11 are compared in a one-way analysis of variance (ANOVA) with Dunnett's multiple comparison–corrected post hoc test against wild-type NK1R stimulated with SP6-11, (^ = p ≤ 0.033, ^^ = p ≤ 0.002, ^^^ = p ≤ 0.0002, ^^^^ = p ≤ 0.0001). NR, no response. ND, not determined.

| | Ligand | $Ca^{2+}$ $pEC_{50}$ | $Ca^{2+}$ $E_{max}$ | cAMP $pEC_{50}$ | cAMP $E_{max}$ | IPone $pEC_{50}$ | IPone Emax |
|---|---|---|---|---|---|---|---|
| NK1R WT | SP | 8.7 ± 0.1 (9) | 72 ± 6 (9) | 6.8 ± 0.1 (12) | 184 ± 24 (12) | 8.0. ± 0.2 (3) | 105 ± 3 (3) |
| | SP6-11 | 9.4 ± 0.1 (8) | 72 ± 4 (8) | 5.4 ± 0.5 (12) | 106 ± 15 (11) | 7.7 ± 0.2 (3) | 101 ± 7 (3) |
| N85D | SP | 8.8 ± 0.1 (4) | 76 ± 3 (4) | 6.4 ± 0.2 (3) | 84 ± 16 (3) | ND | ND |
| | SP6-11 | 8.0 ± 0.2 (3) ^^ | 84 ± 6 (3) | NR | NR | ND | ND |
| N85Q | SP | 8.1 ± 0.4 (5) | 70 ± 4 (5) | 5.9 ± 0.2 (4) * | 68 ± 12 (4) * | ND | ND |
| | SP6-11 | 7.2 ± 0.5 (4) ^^^^ | 66 ± 7 (4) | NR | NR | ND | ND |
| N89D | SP | 8.5 ± 0.2 (5) | 76 ± 2 (5) | 5.0 ± 0.5 (4) **** | 141 ± 38 (4) | ND | ND |
| | SP6-11 | 6.2 ± 0.2 (4) ^^^^ | 96 ± 18 (4) | NR | NR | ND | ND |
| H108A | SP | 8.6 ± 0.1 (5) | 78 ± 3 (5) | 6.1 ± 0.1 (4) | 44 ± 6 (4) ** | ND | ND |
| | SP6-11 | 7.0 ± 0.1 (4) ^^^^ | 81 ± 3 (4) | NR | NR | ND | ND |
| H108Q | SP | 8.7 ± 0.2 (5) | 69 ± 3 (5) | 5.3 ± 0.1 (3) | 115 ± 14 (3) | ND | ND |
| | SP6-11 | 7.9 ± 0.2 (4) ^^^ | 75 ± 6 (4) | 5.3 ± 0.1 (3) | 49 ± 5 (3) | ND | ND |
| Y287F | SP | 8.3 ± 0.3 (5) | 58 ± 2 (5) | 6.3 ± 0.1 (3) | 68 ± 13 (3) * | ND | ND |
| | SP6-11 | 7.7 ± 0.3 (4) ^^^ | 61 ± 5 (4) | NR | NR | ND | ND |

| | | | | | | | |
|---|---|---|---|---|---|---|---|
| Y287H | SP | 8.0 ± 0.2 (5) | 77 ± 3 (5) | 5.1 ± 0.2 (3) **** | 138 ± 70 (3) | ND | ND |
| | SP6-11 | 6.3 ± 0.5 (3) ^^^^ | 82 ± 19 (3) | NR | NR | ND | ND |
| M174I | SP | 8.2 ± 0.2 (5) | 64 ± 7 (5) | 6.4 ± 0.1 (3) | 55 ± 6 (3) * | ND | ND |
| M174V | SP | 8.5 ± 0.2 (5) | 41 ± 2 (5) **** | 6.4 ± 0.1 (3) | 91 ± 12 (3) | ND | ND |
| M181V | SP | 8.4 ± 0.4 (5) | 70 ± 5 (5) | 6.5 ± 0.1 (4) | 185 ± 21 (4) | ND | ND |
| M181F | SP | 8.2 ± 0.2 (5) | 76 ± 5 (5) | 6.5 ± 0.1 (3) | 150 ± 22 (3) | ND | ND |
| R177K | SP | 8.7 ± 0.3 (6) | 76 ± 4 (6) | 6.0 ± 0.1 (4) * | 149 ± 13 (4) | ND | ND |
| R177M | SP | 8.5 ± 0.2 (9) | 75 ± 3 (9) | 5.5 ± 0.2 (8) **** | 62 ± 22 (8) *** | 7.4 ± 0.1 (3) | 107 ± 2 (3) |
